# The complex interplay between microalgae and the microbiome in production raceways

**DOI:** 10.1101/2025.01.20.633910

**Authors:** Judith Traver-Azuara, Caterina R. Giner, Carmen García-Comas, Ana Sánchez-Zurano, Martina Ciardi, Gabriel Acién, Sofiya Bondarenko, Aleix Obiol, Ramon Massana, Maria Montserrat Sala, Ramiro Logares, Pedro Cermeño

**Author notes:** **Corresponding authors:** Judith Traver Azuara Pedro Cermeño Ainsa Ramiro Logares Carmen García-Comas Rubio.

## Abstract

Algae-associated microbiomes are underexplored, limiting our understanding of their influence on the productivity of large-scale microalgae reactors. To address this, we monitored microbial dynamics in two microalgae biomass production raceways over two 8-month intervals inoculated with *Desmodesmus armatus*. One reactor was fed with wastewater, while the other received clean water and fertilizers. Metabarcoding of the 18S and 16S rRNA genes revealed a high microbial diversity across two time series, showing thousands of eukaryotic and prokaryotic species growing alongside the microalgae. Chlorophyta and Fungi were the dominant eukaryotic groups, while Alphaproteobacteria, Gammaproteobacteria, Actinobacteria, and Bacteroidia dominated the prokaryotic communities. We found contrasting ASVs (Amplicon Sequence Variant) patterns between healthy (*D. armatus* abundance >70%) and unhealthy (*D. armatus* abundance <20%) microbiomes, across reactors and time series. Network analysis identified up to 10 potential ecological interactions among *D. armatus* and its microbiome, predominantly positive. Our results suggest a link between microbiome composition and *D. armatus* abundance. Specifically, ASVs associated with a healthy microbiome were positively correlated with *D. armatus*, while ASVs characteristic of an unhealthy microbiome were negatively correlated. Potentially pathogenic bacteria included *Mycobacterium* and *Flavobacterium*, whereas potentially beneficial taxa included *Geminocystis, Thiocapsa, Ahniella* and *Bosea*. Several fungal ASVs showed context-specific associations, whereas specific *P. tribonemae*, *A. parallelum*, *A. desmodesmi*, *Aphelidiomycota* sp., Rozellomycota sp. and, *Rhizophidium* sp. ASVs were identified as potentially harmful. This study reveals the striking diversity and complexity of microalgae-associated microbiomes within raceways, providing valuable insights for optimizing industrial production processes, particularly for wastewater treatment and sustainable green biomass generation.

Graphical Abstract.
General overview of the metabarcoding methodology. Step 1: Sample collection, 2: DNA extraction of the samples, 3: PCR and DNA sequencing of the 18S and 16S rRNA genes, 4: Processing and quality filtering of the DNA sequencing data, 5: Taxonomic, community, and network analyses and, 6: Interpretation and discussion of the results.

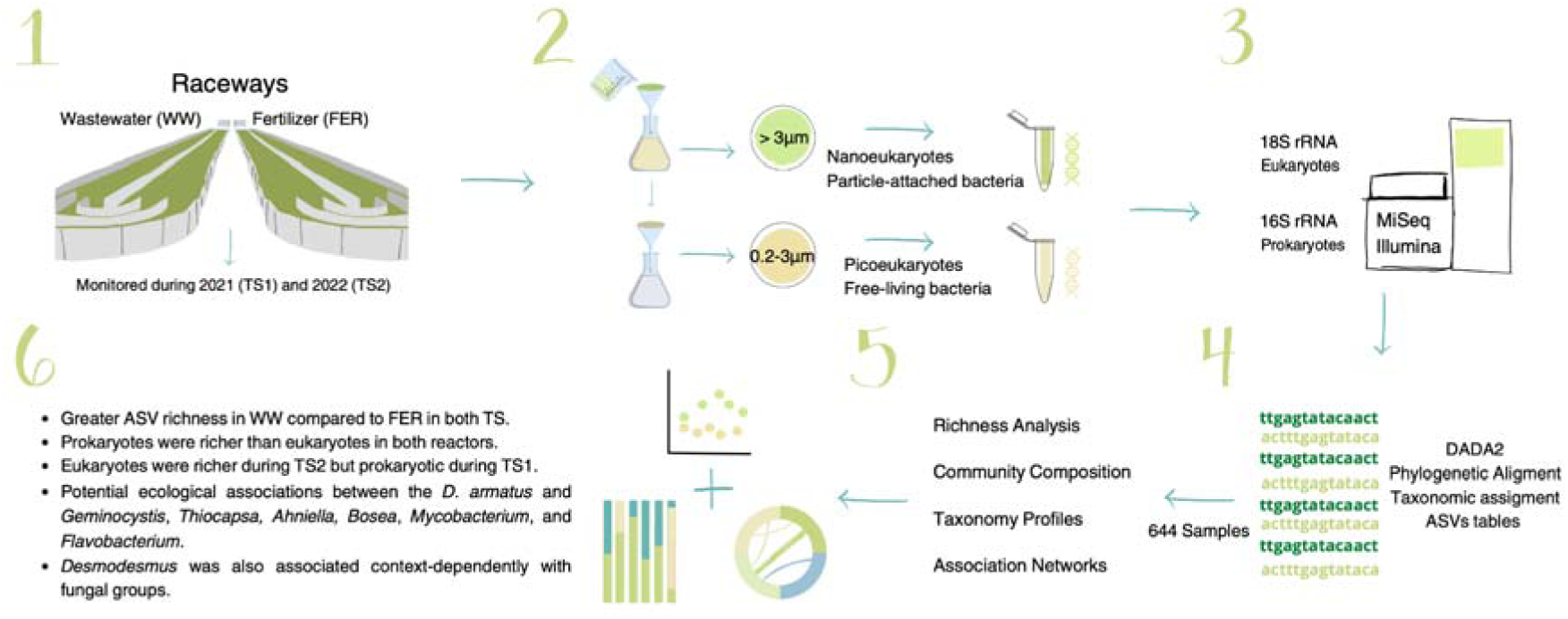

## Introduction

Microalgae are a sustainable alternative to fossil fuels and a possible solution for the world’s food supply (Ahmad & Ashraf, 2023; Kusmayadi et al., 2021; Wijffels & Barbosa, 2010). Their high efficiency in converting sunlight energy and carbon dioxide into valuable organic compounds, coupled with their versatility in producing biofuels, biochemicals, bioplastics, and high-value nutritional supplements, place microalgae at the forefront of sustainable innovation, providing a wealth of opportunities for green industries and circular bioeconomy (Chhandama et al., 2021; D’Alessandro & Antoniosi Filho, 2016; Dolganyuk et al., 2020). Despite the high expectations generated around microalgae during the last two decades, the large-scale implementation of microalgae production systems still faces significant challenges (Borowitzka & Vonshak, 2017; Khan et al., 2018; Novoveská et al., 2023). Among them, understanding and managing the microbiome associated with microalgae stands as a crucial barrier to the large-scale production of microalgae biomass and the commercial expansion of its derivatives (Lian et al., 2018). The concept of microbiome, initially proposed by Whipps and coworkers (1988), includes the microbial community with specific properties and functions and its interactions with the environment (Berg et al., 2020). The microbial community is composed not only of phototrophic organisms, but also of bacteria, protozoa, yeast and fungi (Bani et al., 2020). The composition and dynamics of the microbial community influence the performance of large-scale microalgae production systems (Bani et al., 2020, Lian et al 2018) when the beneficial microbial composition, which maintains the balance in the system, is affected by harmful and pathogenic microorganisms (Zhou et al., 2009) and it results in productivity losses or culture crashes (Molina-Grima et al., 2022).

Currently, available methods for studying microbiomes, such as high-throughput sequencing technologies, are revolutionizing our understanding of microbial ecosystems by determining their taxonomic composition (metabarcoding) and thus revealing their vast diversity and complexity (Berg et al., 2020; Lakaniemi et al., 2012). Recent evidence demonstrates that microbial diversity is essential for maintaining the health and functionality of natural ecosystems (Delgado-Baquerizo et al., 2016; Fierer, 2017; Smith, 2007). A diverse microbial community can adapt to a range of environmental conditions, enhancing the overall ecosystem’s ability to resist and recover from disturbances (Awasthi et al., 2014). In microalgae reactors, certain bacteria and fungi enhance microalgal growth and strengthen the ecosystem’s ability to withstand fluctuations in temperature, pH levels, and salinity (Lian et al., 2018; Seymour et al., 2017). Many microorganisms are pivotal for supporting biodiversity and facilitating key ecosystem processes such as nutrient recycling and oxygenation of water bodies. For example, heterotrophic bacteria, by breaking down organic carbon in microalgae culture systems, release carbon dioxide, vitamin B12, and specific hormones, which microalgae then use to enhance their growth and development (Alam et al., 2022). Furthermore, a diverse microbial community plays a crucial role in defense mechanisms, serving as a natural control (Kim Hue et al., 2018; Zhou et al., 2009). An illustrative example of this is the relationship between the microalgae *Emiliania huxleyi* and the bacteria *Phaeobacter gallaeciensis*, where the bacterium produces antibiotic compounds to shield the host from other bacterial pathogens (Seymour et al., 2017). Also, some heterotrophic bacteria uptake dead microalgae cells from the culture medium and reduce unwanted contaminants (Alam et al., 2022). Specific microbial interactions have proven key in naturally managing biological threats like pathogens, parasites, and competitors (Grube et al., 2015; Ramanan et al., 2016), thus reducing the need for chemical treatments and optimizing culture conditions for enhanced productivity and superior product quality (Jiang et al., 2021). Specific bacteria are capable of degrading pollutants or scavenging heavy metals, which relieves chemical stress on the microalgae while making these microalgal-bacterial consortia ideal for bioremediation purposes (Jiang et al., 2021; Muñoz et al., 2003; Subashchandrabose et al., 2011). Likewise, specific members of the microalgae microbiome significantly influence the macromolecular composition of microalgae, promoting the synthesis of valuable compounds such as lipids (Das et al., 2022) and pigments (Wang & Hong, 2022). For example, in *Chlorella vulgaris* cultures, the lipid and biomass production are enhanced by the presence of the bacteria *Mesorhizobium sangaii* (Wei et al., 2020) while growth and lipid content are promoted by the presence of the fungi *Mucor indicus* (Rodrigues Reis et al., 2018). On the other hand, within the microbial diversity are also found microorganisms that negatively impact the commercial production of microalgae (Day et al., 2017). Their presence can imply grazing, consuming resources in competition with the microalgae cultured, or causing disease (Molina-Grima et al., 2022).

Therefore, optimizing the management of algae-associated microbiomes offers a promising solution to address the scalability challenges in microalgae production systems (Lian et al., 2018). Central to this effort is identifying which microbial species have beneficial or deleterious effects on the microalgae-microbiome community and the performance of the system. Furthermore, monitoring the microbiome for early warning signals – such as changes indicative of ecological imbalances, or dysbiosis – is crucial for anticipating productivity declines. Addressing these questions could unlock more robust, efficient, and ultimately scalable microalgae culture systems. To achieve this objective, a thorough understanding of the diversity, composition, and dynamics of the algal microbiome at industrial scales is required—a topic that remains largely unexplored.

In this study, °the microbial communities in two raceway reactors were monitored over two 8-month periods, in 2021 (TS1) and 2022 (TS2). Samples were collected every 2-3 days to track community dynamics and reconstruct putative interaction networks. The reactors were initially inoculated with the microalgae *Desmodesmus armatus* and operated under different nutrient regimes, with one receiving wastewater (WW) and the other clean water with fertilizers (FER). Microbial communities were analyzed three times per week using metabarcoding of 18S and 16S rRNA genes (ca. 74-88 data points per time series and raceway). The starting hypotheses were: 1) microbial community composition and diversity would differ between the WW and FER reactors due to distinct nutrient inputs, 2) microbial dynamics would follow stable patterns in both reactors and 3) different microorganisms would be associated with healthy and unhealthy microbiomes conditions in the microalgae culture reactors.

## Materials and methods

### Raceway operation and sample collection

Microalgae were cultured in two outdoor raceway reactors located at the Institute of Agricultural and Fisheries Research and Training (IFAPA) in Almería, Spain. Two time series (TS) were produced, TS1 spanned from May 21 to December 3, 2021 and TS2 from April 20 to December 21, 2022. The raceway reactors consisted of a polypropylene algal pond of 50 m length channel (0.46 m high × 1 m wide). The total surface area was 80m^2^ with a volume of 12 m^3^. A rotating paddle wheel was continuously circulating and mixing the culture. The systems were operated under natural day/night light conditions, open-air, and in semi-continuous mode at a dilution rate of 0.20 day^−1^. The fertilized culture (FER) was sustained with agricultural water supplemented with commercial-grade fertilizers at constant nitrate and phosphate concentrations. The proportions were: 0.9 mg NaNO_3_, 0.18 mg MgSO_4_, 0.14 mg KH_2_PO_4_, and 0.02 mg Karentol. In contrast, the wastewater culture (WW) was sustained with primary wastewater from the University of Almería and, when it was limited, by urban primary wastewater from the Almería wastewater treatment plant. The composition of the primary wastewater was variable, fluctuating on a weekly basis.

Raceways were inoculated at 10% of their total volume with *Desmodesmus armatus,* a green microalga from the culture collection of the University of Almería’s Department of Chemical Engineering (Spain). Then, the raceways (Graphical abstract, panel 1) were filled with either primary-treated wastewater or fertilized water until the desired culture depth of 0.15 m was reached. To keep the volume steady, water lost to evaporation was replenished daily.

Samples for community characterization were sampled Monday, Wednesday, and Friday, specifically 30 mL from the FER reactor and 15 mL from the WW reactor. This volume was sequentially vacuum filtered using 47mm polycarbonate filters with nominal pore sizes of 3 μm and 0.2 μm to separate microorganisms into nano- and picoplankton size classes, respectively. Then, those microbial samples were immediately stored at -80°C and sent to the laboratory at the Institute of Marine Sciences (ICM), CSIC in Barcelona for metabarcoding analysis.

### Analytical Methods

Biomass and physicochemical variables were sampled daily from Monday to Friday. Biomass concentration was determined by dry weight after filtering 100 mL of the culture through Macherey–Nagel glass fiber filters (MN 85/90, Thermo Fisher Scientific, Spain). Then, samples were dried at 80°C for 24 hours. The biomass consisted of a mix of microalgae and its microbiome. Nitrate (N-NO□□) levels were quantified using a GENESYS 10S UV–Vis spectrophotometer (Thermo Fisher Scientific, Spain) at wavelengths between 220 and 275 nm. Ammonium (N-NH□□) concentration was determined using the Nessler reagent method. Nutrient removal efficiency was assessed by comparing the concentration of inorganic nutrient input with the output. Biomass, nitrogen concentrations, and nitrogen removal percentages are displayed in Tables S1 and S2.

### DNA extraction, sequencing, and processing

Community DNA was extracted directly from polycarbonate filters, using DNeasy PowerSoil Pro Kit (Qiagen) following the manufacturer’s instructions, and quantified with NanoDrop One/OneC UV-Vis (Thermo Scientific™). The amount of total DNA recovered from samples ranged from 1 to ca. 300 ng/μL.

The DNA samples were normalized according to their DNA concentration to be used as templates to prepare 16S rRNA (region V4-V5, (Parada et al., 2016)) and 18S rRNA (region V4, (Balzano et al., 2015)) libraries. Both regions 16S rRNA (V4-V5) and 18S rRNA (V4) were amplified with the specific primers: 16S-V4-V5 rRNA (515F-Y (5’-GTGYCAGCMGCCGCGGTAA); 926R (5’-CCGYCAATTYMTTTRAGTTT)) and 18S -V4 (V4F (5’-CCA GCA SCY GCG GTA ATT CC-3’); V4RB (5’-ACT TTC GTT CTT GAT YRR-3’)).

PCRs and library preparation protocols are detailed in Method S1. Once the libraries were normalized, they were prepared for subsequent sequencing. Sequencing was performed on Illumina’s MiSeq with 2×300 bp reads using v3 chemistry. 16S and 18S rRNA amplicons were sequenced in different runs, and a total of 10 sequencing runs were used to sequence the entire dataset. PCRs, library preparation, and sequencing were carried out at the sequencing service of the Centre for Genomic Regulation (CRG) of the Biomedical Research Park of Barcelona (PRBB) in Barcelona, Spain.

Raw sequences were initially processed using Cutadapt version 1.16 (Martin, 2011), removing forward and reverse primers. Sequences where both primers could not be identified were removed. Then, we applied the state-of-the-art tool DADA2 (Callahan et al., 2016). Initially, sequences were filtered and trimmed using the function filterAndTrim from the dada2 package. For both the 16S and 18S rRNA data, the filtering parameters ensured high-quality sequence data by trimming low-quality bases (truncLen=c (250,200)), setting a low *Phred* quality threshold for truncation (truncQ=2), limiting the maximum number of expected errors (maxEE=c(2,2)), and removing any sequences with ambiguous nucleotides (maxN=0). Additionally, *PhiX* sequences were removed from the dataset (rm.phix=TRUE). Finally, chimeric sequences were removed using the function *removeBimeraDenovo* with a consensus method, and identical sequences with varying lengths were collapsed using the function *collapseNoMismatch,* setting a minimum overlap of 20 nucleotides. Then, we generated Amplicon Sequence Variants (ASVs). ASV’s taxonomy was assigned using the approach *assignTaxonomy* in DADA2 and different databases. For the 18S dataset, we used the PR2 database version 5.0.0 (Guillou et al., 2012). To improve the eukaryotic taxonomic classification, we compared the taxonomic assignment of the eukaryotic ASVs with the assignments based on the EukV4 reference database (Obiol et al., 2020), which is continuously updated. For the 16S dataset, we used *assignTaxonomy* and then added species information with the addSpecies function, using the SILVA database version 138 (Quast et al., 2012).

Then, an abundance filtering was performed, and all the samples had > 2,000 reads. To improve the quality of the 18S and 16S datasets, a Phylogenetic Alignment of the ASVs was carried out using Mothur (Schloss et al., 2009) version 1.40.5. We visualized the completeness of conserved and variable regions of the aligned ASVs with SeaView version 5.0.5 (Gouy et al., 2021). Those ASVs that did not align properly were removed. A phylogenetic tree was constructed using FastTreeMP v2.1.10 with the parameters -gtr, - gamma, and -nt. These parameters specified the generalized time-reversible (GTR) model and a gamma-distributed shape parameter that facilitates modeling rate variation across nucleotide positions through a discrete gamma distribution with 20 categories, thereby improving the accuracy of phylogenetic likelihood calculations. We removed from the tree those ASVs producing long or isolated branches, which would point to chimeras or sequencing errors. We additionally checked for the presence of chimeras by comparing the ASVs against different databases (NCBI version 2019-02-02, version 2024 downloaded 2024-03-22, SILVA version 138, PR2 version 5.0.0, and Eukv4 version 8). Chloroplast and mitochondria ASVs were removed from the prokaryotic dataset and metazoa and streptophyta were removed from the eukaryotic dataset. The initial eukaryotic dataset that contained all the samples (TS1 and TS2) had 20,836 ASVs and after the filtering steps, we kept 1,778 ASVs (75.73% of the the dataset). In turn, the prokaryotic dataset with all the samples initially had 29,316 ASVs and 9,075 ASVs were kept after filtering which represent 64.96% of the data.

The number of reads per sample was normalized with the *rrarefy* function from the R-package vegan 2.6.4 (Oksanen et al., 2022). We ran 100 rarefactions per sample, with a subsample size of 18,000 reads per sample.

The final dataset contained a total number of 644 samples, 296 from TS1 (148 from each WW and FER reactor) and 348 from TS2 (172 from the WW reactor and 176 from the FER reactor).

### Diversity measurements and community analyses

Richness and community structure analyses were calculated for eukaryotes and prokaryotes. We considered two filter size fractions for prokaryotes and eukaryotes, distinguishing between picoeukaryotes (0.2-3 µm) and nanoeukaryotes (> 3 µm), as well as free-living (0.2-3 µm) and particle/protist-attached bacteria (> 3µm). For each sample, we calculated the richness as the number of ASVs per sample. Beta-diversity between samples was estimated using Bray-Curtis dissimilarities.

Taxonomic profiles were depicted at the Class level, highlighting the 10 most abundant groups, with the remaining rare groups categorized as “Other”.

To determine which ASVs consistently and most likely explained differences between contrasting healthy and unhealthy microbiomes in the raceways, we classified samples as healthy when the *D. armatus* relative abundance was >70% of the total abundance of eukaryotes (all size fractions) and as unhealthy when the *D. armatus* abundance was between 20% and 10%. We then applied a Linear discriminant analysis Effect Size (LEfSe) (Segata et al., 2011), which evaluates the effect size and statistical significance of each ASV (eukaryotic and prokaryotic) for the two contrasting groups. For the WW reactor, we identified 16 samples with healthy microbiomes and 14 samples with unhealthy ones from TS1, and 13 healthy and 17 unhealthy from TS2. For the FER reactor, we obtained 17 healthy and 22 unhealthy microbiomes from TS1 and 22 healthy and 14 unhealthy microbiomes from TS2 (see Method S2). ASVs with Linear discriminant analysis (LDA) scores ≥2.0 and p-values <0.05 were identified as the most significantly different between groups, and healthy and unhealthy samples were characterized by their top-five most representative ASVs (Table 2 and 3). The analysis was performed on the community composition with D. armatus included.

Graphs were generated with R software v.4.1.3 (Team, 2022) in RStudio 2022.07.2+576 (Team, 2022) with the packages Vegan 2.6.4 (Oksanen et al., 2022), ggplot2 v.3.5.1 (Wickham et al., 2016), tidyverse v.1.3.1 (Wickham, 2021), dplyr v.1.1.4 (Wickham et al., 2023), zoo v.1.8-12 (Zeileis et al., 2023), reshape2 v.1.4.4 (Wickham, 2020), gridExtra v.2.3 (Auguie & Antonov, 2017) and LefSe v.1.1.2 (Segata et al., 2011).

### Network analyses

To determine relationships between pairs of ASVs, we constructed microbial co-occurrence networks with Flashweave (Tackmann et al., 2019). Flashweave integrates microbial relative abundance data with environmental metadata to predict associations representing potential ecological interactions between ASVs. It is based on a flexible Probabilistic Graphical Model that allows inferring microbial association networks based on co-occurrence (species frequently appearing together) or co-exclusion (species rarely appearing together) of taxa across samples while controlling for environmental context-dependency. The result is an ecological network where nodes represent microbial species (ASVs) and edges indicate potential positive (co-occurrence) or negative (co-exclusion) interactions between them. As in previous work (Deutschmann et al., 2024), to normalize the data we applied a centered-log-ratio (clr) transformation separately to the bacterial and eukaryotic abundance tables (that combine pico- and nano-size fractions) before merging them. This transformation was applied to the non-rarefied ASV tables. In Flashweave, we set the parameter sensitive=true, to increase the sensitivity of the algorithm, allowing for the detection of weaker associations between taxa to the detriment of false positives, which we then trimmed through association strength filtering. We set the parameter heterogeneous=True to account for environmental context-dependency, that is, allowing associations to vary according to the environment (Tables S7-S10). From the output network (Tables S11 and S12), the ASV-environment associations were removed, and we obtained the initial network (Tables S13 and S14). This initial network had more than 3,000 nodes and 8,000 edges and was filtered by the weight and the co-occurrence of the associations. On the one hand, from the initial network we filtered associations by their weight > |±0.5| and by the ASVs co-occurrence (> 5% co-occurrence= > in 8 samples) to obtain the filtered network (Table S15 and S16). On the other hand, from the initial network we selected the associations involving *D. armatus* ASVs and filtered them by > 4% co-occurrence (> 7 samples) to obtain the *Desmodesmus*-microbiome network (Tables S17 and S18). Flashweave was run in Julia v.1.9.1 (https://julialang.org/ 2023-06-07).

To characterize the network structure, we considered sixteen network metrics: 1) number of nodes; 2) number of edges; number of 3) positive and 4) negative edges; percentage of 5) positive and 6) negative edges; 7) the average strength of positive associations; 8) edge density to indicate network connectivity as the number of actual edges divided by the number of all possible edges; 9) average path length to indicate distance between nodes as the average length of all shortest paths between pairs of nodes in a network; 10) transitivity to evaluate clustering within the network which is the probability that the nodes’ neighbors are connected; 11) the diameter to know the longest distance between two nodes via the shortest path, estimated by finding the shortest paths between all pairs of nodes and identifying the longest one; 12) the average degree as the mean number of connections (degrees) per node; 13) degree centralization to measure how much the network relies on a few highly connected nodes, calculated by comparing the degree of each node to the maximum degree in the network and summing these differences; 14) assortativity (degree) to determine if nodes with similar degree tend to connect; indicating if well-connected ASVs are likely to interact with other highly connected ASVs. Similarly, 15) assortativity (Euk-Prok) is positive if eukaryotic organisms, such as microalgae, preferentially connect with other eukaryotes, while prokaryotic organisms tend to associate with their own kind, highlighting the distinct community structures and interactions between these groups; lastly, we computed the 16) modularity to know the degree to which a network can be divided into distinct subcommunities, groups, or clusters. Its values can range from -1 to 1, where a positive value indicates the presence of well-defined sub-communities or tight-knit sub-groups in the culture; organisms within those sub-communities interact more frequently with each other than with others from the community.

Network metrics were computed with the igraph v.1.4.3 R-package (Csárdi et al., 2023), and associations were visualized with the circlize v.0.15 R-package (Gu, 2022).

## Results

### Biomass, nitrogen consumption, and nutrient uptake

In both reactors and throughout both time series, nutrient concentrations varied significantly, which resulted in different biomass yields (Table S1 and S2). Total biomass varied over time in both the WW and FER raceway reactors across TS1 and TS2 (Figure 1a). In TS1, the highest biomass concentration in the WW reactor was observed in October, reaching approximately 0.8 g/L. Conversely, the peak for the FER reactor was in July, slightly below 1 g/L. The WW reactor displayed more variability in biomass concentration than the FER reactor in both time series (TS1: F = 1.40, p-value > 0.02, and in TS2: F = 3.64, p-value < 2.2e-16). This resulted from the large fluctuations in nutrient concentrations found in the wastewater, which often exceeded the constant levels maintained in the FER reactor (Figure 1b, c). In TS2, the biomass concentration was significantly higher in the WW reactor compared to the FER (W = 47940, p-value < 2.2e-16; Figure 1a), which matches greater nutrient inputs in the WW reactor (W = 50301, p-value < 2.2e-16; Figure 1b). In TS2, the maximum biomass concentration in the WW reactor was 1.9 g/L in August, whereas the FER peaked 0.96 g/L in May.

**Figure 1.**
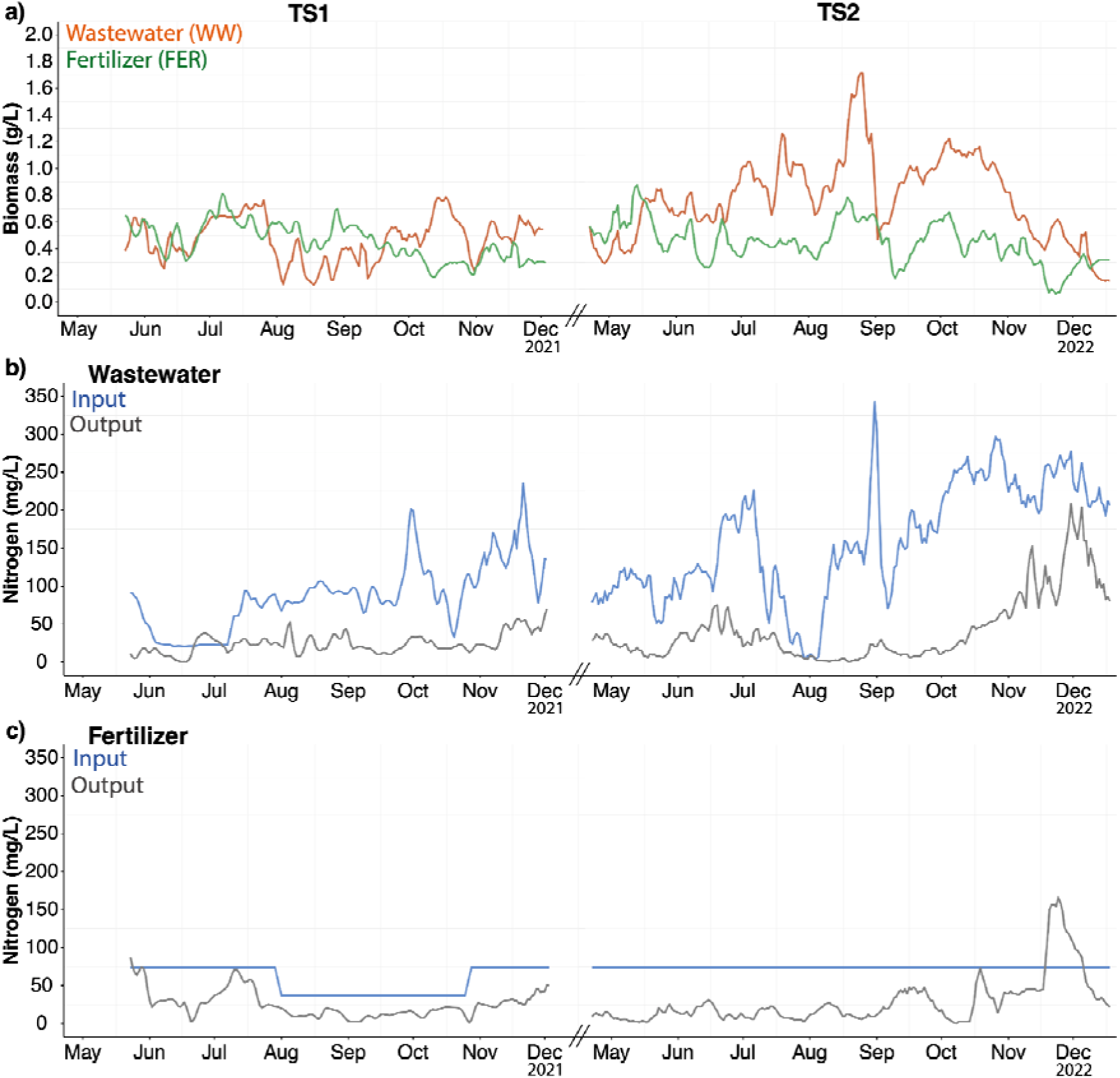
Biomass and nutrient removal efficiency. Biomass (a) and nitrogen (b, c) concentration (moving average with a 3-day window) in the wastewater and fertilizer reactors during TS1 (from May to December 2021) and TS2 (from April to December 2022). In the wastewater reactor, nitrogen represents N-NH_4_^+^ for the input, whereas NO_3_ and NH_4_ for the output. In the fertilizer treatment, nitrogen represents NO_3_.

Inorganic nitrogen (N) input to the reactors showed significantly greater variability in WW than in FER in both TS (TS1: F = 6.32, p-value < 2.2e-16, TS2: F = Inf, p-value < 2.2e-16), and almost all nitrogen was removed from the reactors (Figure 2b and c). During both time series, the ammonium concentration in the outputs was significantly lower than in the inputs (p < 0.05). With nitrogen removal efficiencies above 90% in 16% and 14% of cases in WW and FER raceways, respectively.

**Figure 2.**
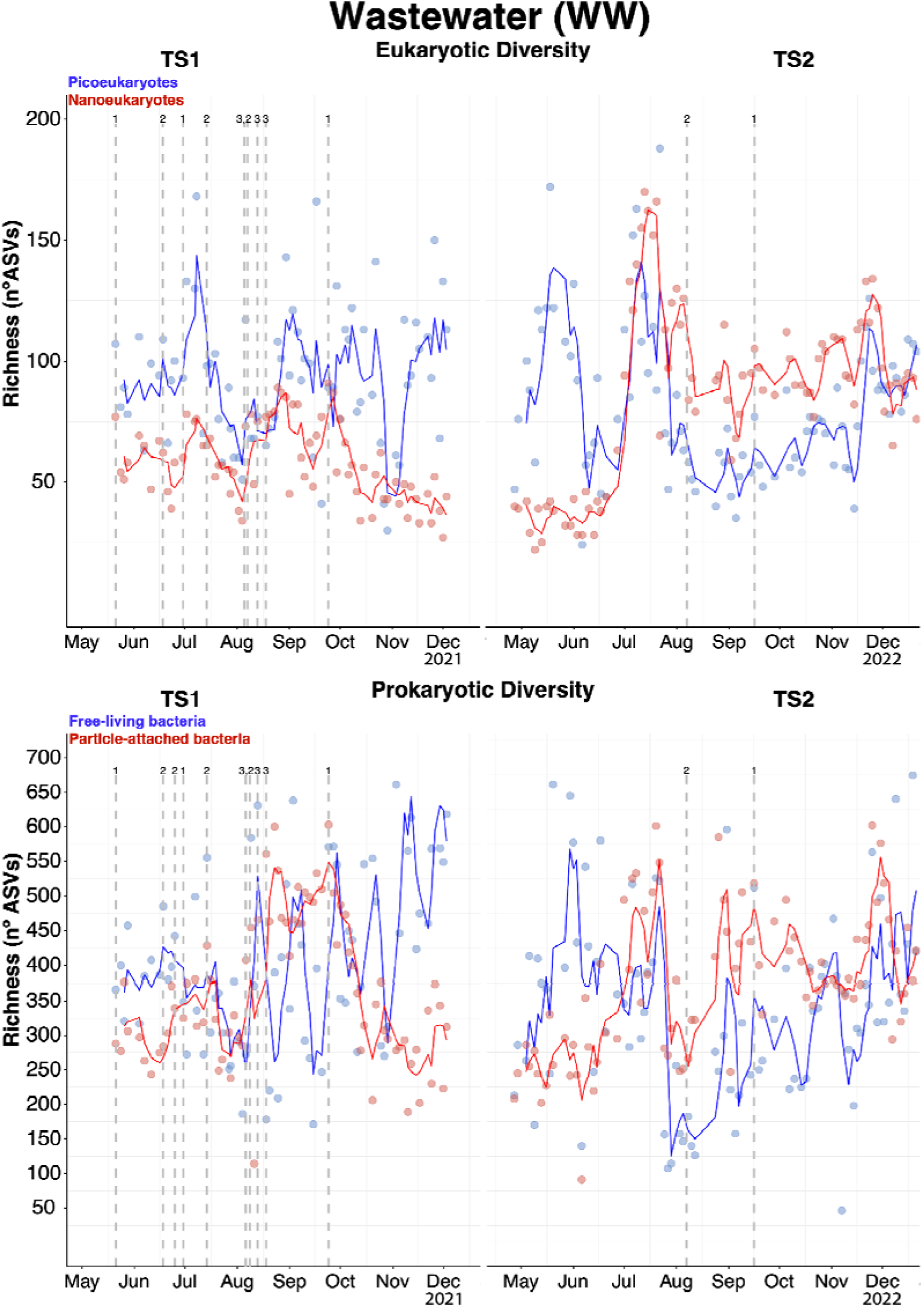
Microbial ASV richness in the wastewater reactor over two independent time series, TS1 and TS2. Each dot represents the richness (number of ASVs) in a single sample. The line shows a 3-point moving average of these richness values to illustrate the overall trend. The dashed lines indicate a change in the water type entering the raceway: 1) university wastewater, 2) city wastewater, and 3) city wastewater plus fertilizer culture or fertilizers.

### Microbial community richness within the reactors

ASV richness fluctuated significantly, spanning up to 2-3 times its minimum values along the time series. In the WW reactor, we observed a summer (July-August) decline in both eukaryote and prokaryote richness across size fractions (Figure 2). This drop was stronger in the pico size fraction than in the nano size fraction. Indeed, the pico size fraction showed stronger fluctuations than the nano fraction. In the FER reactor, we did not find any temporal pattern (Figure 3).

**Figure 3.**
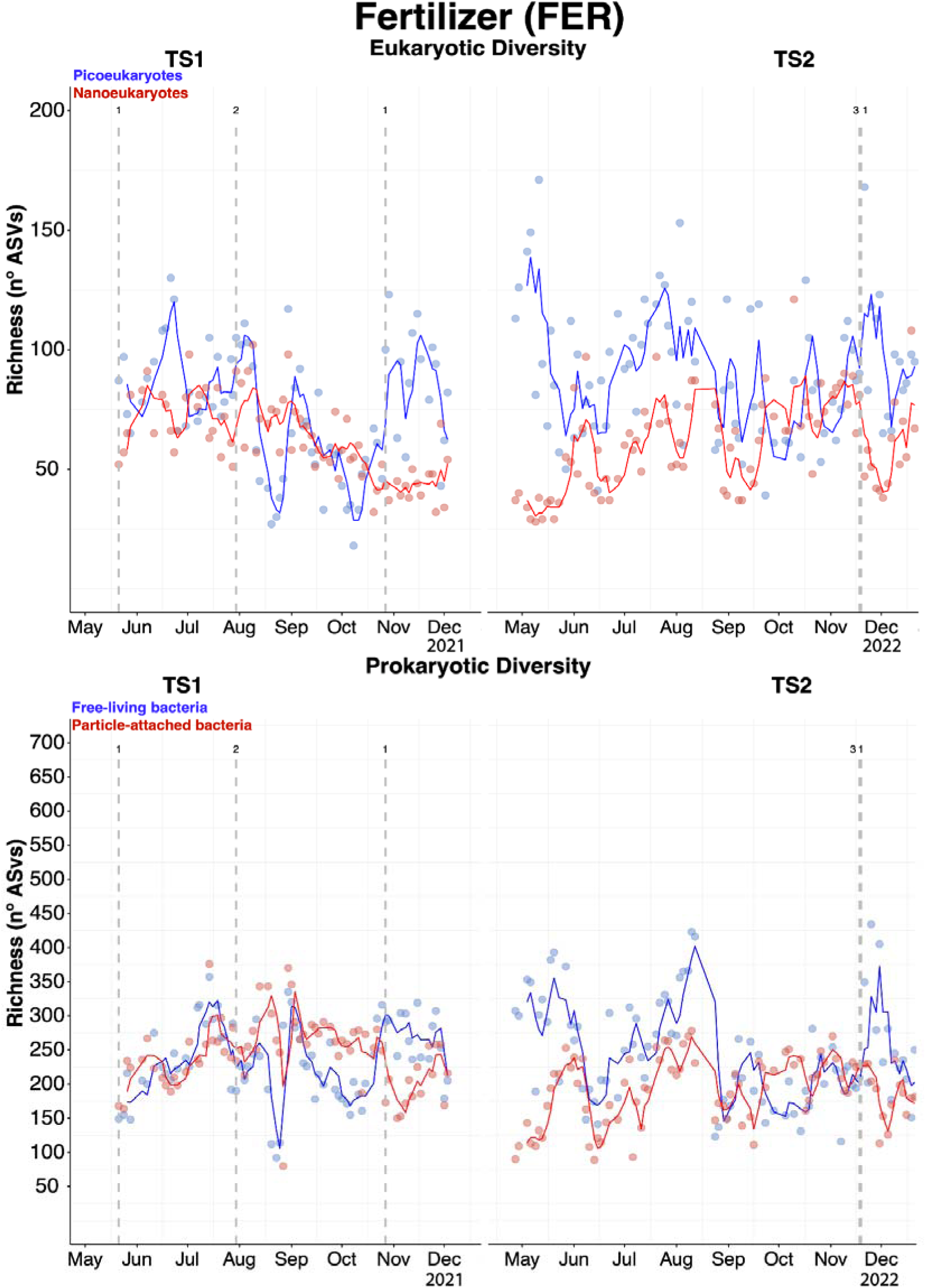
Microbial ASV richness in the fertilizer reactor over two independent time series, TS1 and TS2. Each dot represents the richness (number of ASVs) in a single sample. The line shows a 3-point moving average of these richness values to illustrate the overall trend. The dashed lines indicate a change in the nutrient concentration of the water entering the raceway: 1) 50%, 2) 25%, and 3) 100%.

Both eukaryotes and prokaryotes showed greater ASV richness in the WW than in the FER reactor for both TS1 and TS2 (Figure S1 a, b). Furthermore, in both reactors, the prokaryotic compartment was significantly richer than the eukaryotic compartment (Figure S1 c, d). Finally, the eukaryotic compartment was significantly richer in both WW and FER reactors during TS2 but prokaryotic was significantly richer in TS2 (Figure S1 e, f).

In terms of total number of ASVs, we found approximately 4 times more prokaryotic ASVs than eukaryotic ASVs in the WW reactors, and about 3 times more prokaryotes than eukaryotes in the FER reactors (Table 1). Among the eukaryotes, at the phylum level, just a few (on average 8% in TS1 and 5% in TS2) corresponded to Chlorophyta. Around 1/3 of the eukaryotic ASVs corresponded to fungi (Table 1).

**Table 1.** Number of eukaryotic and prokaryotic ASVs in WW and FER raceway reactors across TS1 and TS2 and the distinction between Chlorophyta, Fungi, and Other Eukaryota. *Other Euks*. category corresponds to eukaryotic ASVs, which are not Chlorophyta or Fungi. In brackets is the percentage of ASVs out of the total number of eukaryotic ASVs.

| TS Reactor |  | Prokaryotic<br>ASVs | Eukaryotic<br>ASVs | Chlorophyta<br>ASVs | Fungi<br>ASVs | Other Euks.<br>ASVs |
| --- | --- | --- | --- | --- | --- | --- |
| TS1 | WW | 4541 | 685 | 52 (7.6%) | 266 (38.8%) | 367 (53.6%) |
|  | FER | 2595 | 606 | 50 (8.2%) | 223 (36.8%) | 333 (54.9%) |
| TS2 | WW | 5113 | 1187 | 58 (4.8%) | 432 (36.4%) | 697 (58.7%) |
|  | FER | 2819 | 1043 | 68 (6.5%) | 370 (35.5%) | 605 (58.0%) |

### Microbial community structure and dynamics

Microbial community structure varied significantly across taxonomic groups, size fractions, and time series. Bray-Curtis dissimilarity values ranged from 0 (identical communities) to 1 (completely different communities) (Figures S2a-h, S3a-h), indicating different community turnover rates. As a rule of thumb, we set a low (<0.75) and high (> 0.75) dissimilarity threshold, and calculate the percentage of values above and below this limit for each reactor and TS.

Regarding the picoeukaryotic community, the WW reactor showed a higher percentage of dissimilarity values (>0.75) in TS2 compared to TS1 (81.15% vs 74.50%, X-squared = 81.446, df = 1, p-value < 2.2e-16) (Figure S2i, k). The FER reactor also showed higher dissimilarity values in TS2 than in TS1 (81.40% vs 72.68%, X-squared = 140.81, df = 1, p-value < 2.2e-16). A comparison of both reactors revealed a higher proportion in dissimilarities in WW compared to FER in TS1 (X-squared = 4.605, df = 1, p-value = 0.03188).

Nanoeukaryotic communities in WW and FER reactors showed high resemblance, with dissimilarity values consistently below 0.1 (Figures S2e-h, S3e-h). In the WW reactor, the nanoeukaryotic community composition showed a higher proportion of low dissimilarity values (<0.75) in TS1 than in TS2 (100% vs 46.65%, X-squared = 4210.7, df = 1, p-value < 2.2e-16). However, in the FER reactor, TS2 showed a significantly higher proportion of low dissimilarity values compared to TS1 (73.47% vs 64.2%, X-squared = 129.9, df = 1, p-value

< 2.2e-16). A comparison of the two reactors showed a higher percentage of nanoeukaryotic communities with low dissimilarity in FER compared to WW only in TS2 (X-squared = 1137.1, df = 1, p-value < 2.2e-16).

Free-living bacterial communities showed similar patterns in the WW and FER reactors. In the WW reactor, community dissimilarity was higher in TS2 (85.26%) than in TS1 (75.75%), indicating increased variability over time (X-squared = 186.04, df = 1, p-value < 2.2e-16) (Figure S2j, l). The FER reactor also exhibited greater dissimilarity in TS2 compared to TS1 (77.87% vs 59.46%, X-squared = 186.04, df = 1, p-value < 2.2e-16). A comparison between reactors revealed that WW had a higher percentage of dissimilar communities than FER both time series (TS1: 75.75% vs 59.46%, X-squared = 330.98, df = 1, p-value < 2.2e-16; TS2: 85.26% vs 77.87%, X-squared = 1095.3, df = 1, p-value < 2.2e-16).

Particle-attached bacterial communities showed low dissimilarity values (<0.75) in TS1 compared to TS2 in both reactors, WW reactor (98.87% vs 37.24%, X-squared = 5151.1, df = 1, p-value < 2.2e-16) and FER reactor (77.90% vs 35.77%, X-squared = 2285.6, df = 1, p-value < 2.2e-16) (Figures S2j, l and S3j, l). As before, the comparison between reactors revealed that the WW reactor had a higher percentage of communities with low dissimilarity in both time series but it was significantly different only in TS1 (8.87% v 77.90%, X-squared = 1170.2, df = 1, p-value < 2.2e-16.

The taxonomic composition of the bacterial communities varied significantly between reactors (Figures 4, 5), with over 75% of communities differing between both TS (TS1: X-squared = 367.95, df = 1, p-value < 2.2e-16; TS2: X-squared =490.08, df = 1, p-value < 2.2e-16). Gammaproteobacteria, Alphaproteobacteria, Bacteroidia, Cyanobacteria, and Actinobacteria were prevalent across both WW and FER reactors and throughout TS1 and TS2. Gammaproteobacteria were particularly abundant in the WW reactor (Figure 4), while Alphaproteobacteria thrived in the FER reactor (Figure 5 and Table S19). Beyond this general pattern, there were no consistent taxonomic trends other than specific taxa appearing and disappearing in a seemingly unpredictable manner.

**Figure 4.**
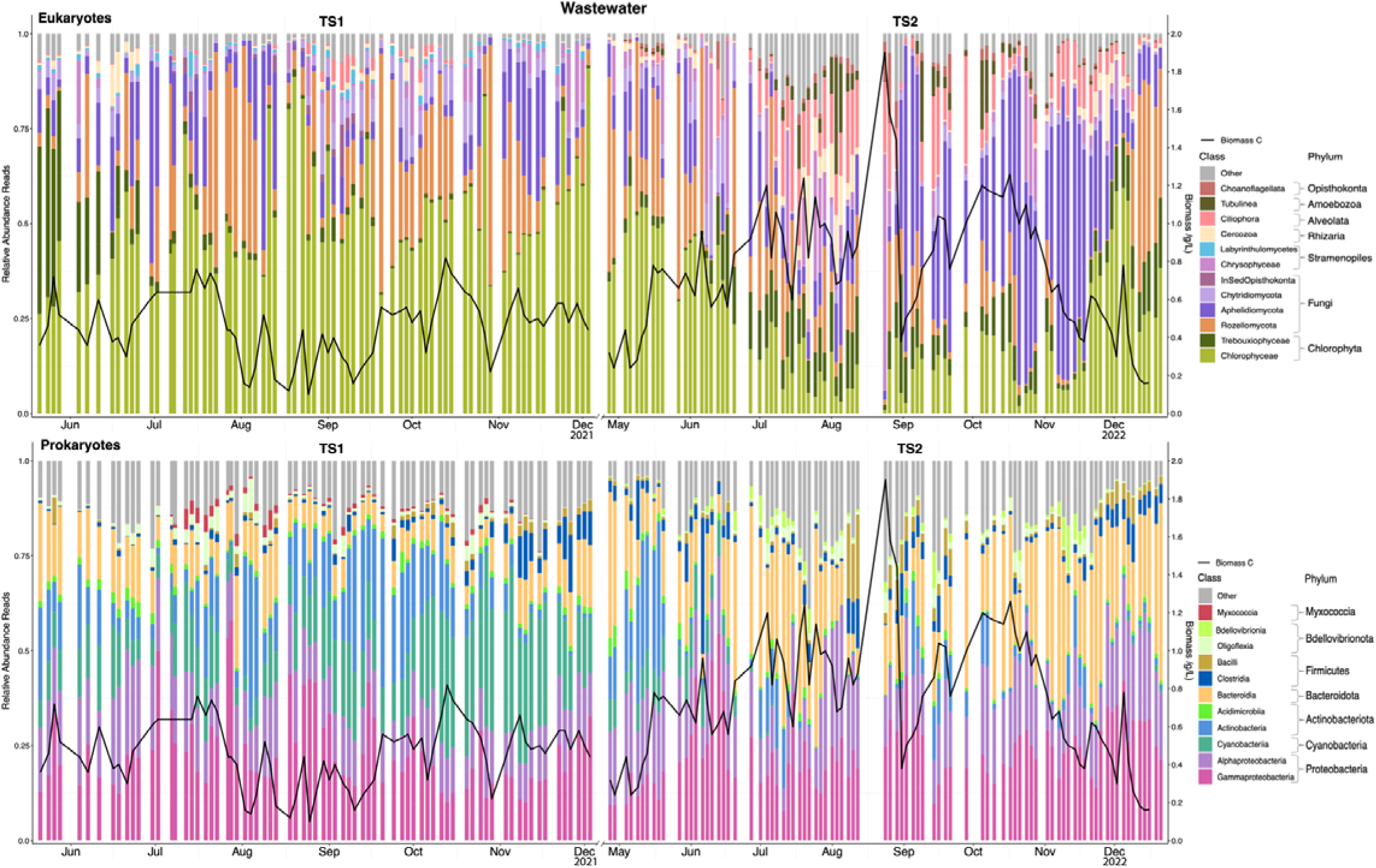
Taxonomic profiles of the microbial communities in the wastewater reactor. Taxonomic composition of eukaryotic (upper panel) and prokaryotic communities (lower panel) over two time series, TS1 (from May to December 2021) and TS2 (from April to December 2022). Taxonomic groups are ranked from bottom to top in decreasing order of abundance, showcasing the ten most abundant classes. The remaining taxa are grouped into the "Other" category (grey bars). The black line corresponds to the biomass in the system.

**Figure 5.**
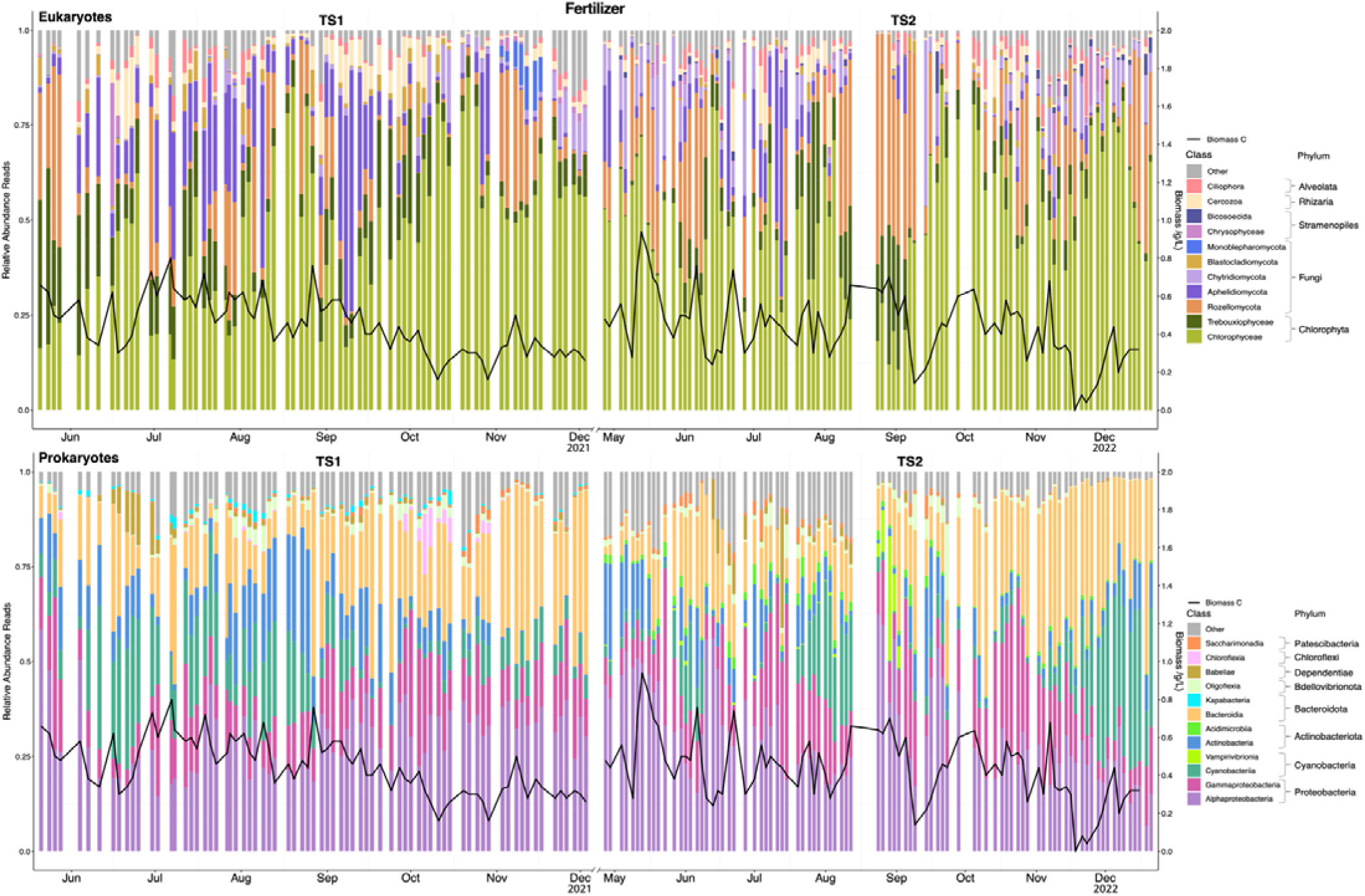
Taxonomic profiles of the microbial communities in the fertilizer reactor. Taxonomic composition of eukaryotic (upper panel) and prokaryotic communities (lower panel) over two time series, TS1 (from May to December 2021) and TS2 (from April to December 2022). Taxonomic groups are ranked from bottom to top in decreasing order of abundance, showcasing the ten most abundant classes. Ther remaining taxa are grouped into the "Other" category (grey bars). The black line corresponds to the biomass in the system.

According to the LEfSe analysis that classifies microbiomes as healthy or unhealthy, the most prevalent ASVs in healthy versus unhealthy microbiomes differed for both reactors and time series except for a few common patterns (Tables 2 and 3). Regarding the top prokaryote component, the cyanobacteria *Geminocystis* PCC-6308 (ASV_2) scored as the top taxon in the healthy microbiomes for the WW reactor in both time series. *Geminocystis* was also found as healthy in FER-TS2 with a different ASV (ASV_1682). On the other hand, ASVs of Actinobacteria *Mycobacterium* were found to be one of the most prevalent in unhealthy microbiomes for both reactors. Its ASV_5 appeared in WW-TS1 and FER-TS1 (Table 2), so the same organism was present in both reactors during this period. Then, ASV_16 appeared only in WW-TS2, while ASV_40 appeared in WW-TS1 and FER-TS2 (Tables S3 and S4). Three different ASVs assigned to *Mycobacterium* suggested it was also present in both time series with small genetic variations. Conversely, some ASVs were not consistently present across both reactors and time series. These microorganisms varied in their presence depending on the health of the microbiome, the reactor type (WW or FER), and the time series (Tables S3, S4, S5 and S6). For example, the Gammaproteobacteria *Thiocapsa* is associated only with healthy microbiomes in WW (TS1 and TS2) while the Alphaproteobacteria *Bosea* in FER (TS1 and TS2) and WW (only TS2); or even the Bacteroidia *Flavobacterium* is associated with both healthy and unhealthy microbiomes in FER and WW reactors during TS1 and TS2 through different ASVs.

**Table 2.**
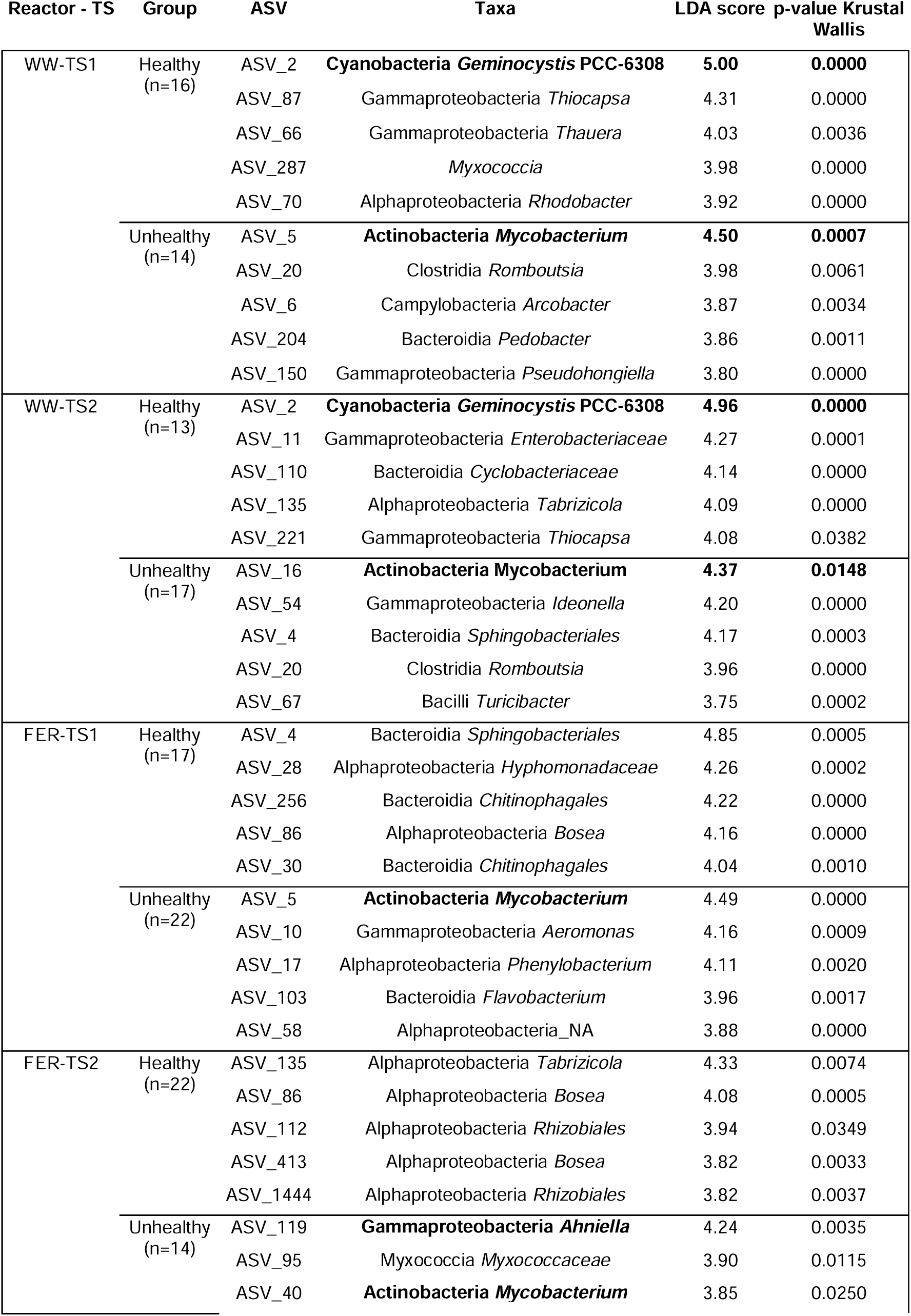
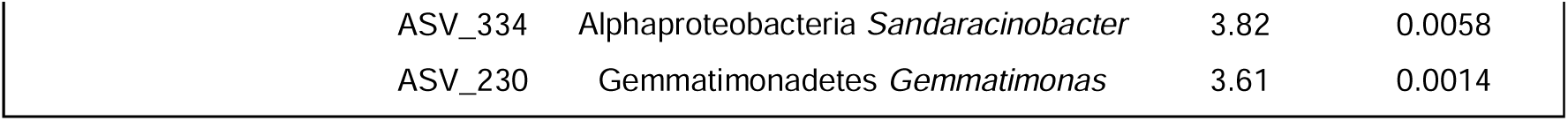
Top-five prokaryotic ASVs related to healthy and unhealthy microbiomes. Healthy (i.e., samples in which *D. armatus* ’s relative abundance was >70% of the total abundance of eukaryotes) and unhealthy (i.e., samples in which *D. armatus* ’s relative abundance was below 20%) microbiome according to LEfSe analysis (see Methods). Bold text corresponds to organisms also found in a positive or negative association with the microalgae in network analyses.

| Reactor - TS | Group | ASV | Taxa | LDA score | p-value Krustal Wallis |
| --- | --- | --- | --- | --- | --- |
| WW-TS1 | Healthy (n=16) | ASV_2 | **Cyanobacteria Geminocystis PCC-6308** | **5.00** | **0.0000** |
|  |  | ASV_87 | Gammaproteobacteria *Thiocapsa* | 4.31 | 0.0000 |
|  |  | ASV_66 | Gammaproteobacteria *Thauera* | 4.03 | 0.0036 |
|  |  | ASV_287 | *Myxococcia* | 3.98 | 0.0000 |
|  |  | ASV_70 | Alphaproteobacteria *Rhodobacter* | 3.92 | 0.0000 |
|  | Unhealthy (n=14) | ASV_5 | **Actinobacteria Mycobacterium** | **4.50** | **0.0007** |
|  |  | ASV_20 | Clostridia *Romboutsia* | 3.98 | 0.0061 |
|  |  | ASV_6 | Campylobacteria *Arcobacter* | 3.87 | 0.0034 |
|  |  | ASV_204 | Bacteroidia *Pedobacter* | 3.86 | 0.0011 |
|  |  | ASV_150 | Gammaproteobacteria *Pseudohongiella* | 3.80 | 0.0000 |
| WW-TS2 | Healthy (n=13) | ASV_2 | **Cyanobacteria Geminocystis PCC-6308** | **4.96** | **0.0000** |
|  |  | ASV_11 | Gammaproteobacteria *Enterobacteriaceae* | 4.27 | 0.0001 |
|  |  | ASV_110 | Bacteroidia *Cyclobacteriaceae* | 4.14 | 0.0000 |
|  |  | ASV_135 | Alphaproteobacteria *Tabrizicola* | 4.09 | 0.0000 |
|  |  | ASV_221 | Gammaproteobacteria *Thiocapsa* | 4.08 | 0.0382 |
|  | Unhealthy (n=17) | ASV_16 | **Actinobacteria Mycobacterium** | **4.37** | **0.0148** |
|  |  | ASV_54 | Gammaproteobacteria *Ideonella* | 4.20 | 0.0000 |
|  |  | ASV_4 | Bacteroidia *Sphingobacteriales* | 4.17 | 0.0003 |
|  |  | ASV_20 | Clostridia *Romboutsia* | 3.96 | 0.0000 |
|  |  | ASV_67 | Bacilli *Turicibacter* | 3.75 | 0.0002 |
| FER-TS1 | Healthy (n=17) | ASV_4 | Bacteroidia *Sphingobacteriales* | 4.85 | 0.0005 |
|  |  | ASV_28 | Alphaproteobacteria *Hyphomonadaceae* | 4.26 | 0.0002 |
|  |  | ASV_256 | Bacteroidia *Chitinophagales* | 4.22 | 0.0000 |
|  |  | ASV_86 | Alphaproteobacteria *Bosea* | 4.16 | 0.0000 |
|  |  | ASV_30 | Bacteroidia *Chitinophagales* | 4.04 | 0.0010 |
|  | Unhealthy (n=22) | ASV_5 | **Actinobacteria Mycobacterium** | 4.49 | 0.0000 |
|  |  | ASV_10 | Gammaproteobacteria *Aeromonas* | 4.16 | 0.0009 |
|  |  | ASV_17 | Alphaproteobacteria *Phenylobacterium* | 4.11 | 0.0020 |
|  |  | ASV_103 | Bacteroidia *Flavobacterium* | 3.96 | 0.0017 |
|  |  | ASV_58 | Alphaproteobacteria_NA | 3.88 | 0.0000 |
| FER-TS2 | Healthy (n=22) | ASV_135 | Alphaproteobacteria *Tabrizicola* | 4.33 | 0.0074 |
|  |  | ASV_86 | Alphaproteobacteria *Bosea* | 4.08 | 0.0005 |
|  |  | ASV_112 | Alphaproteobacteria *Rhizobiales* | 3.94 | 0.0349 |
|  |  | ASV_413 | Alphaproteobacteria *Bosea* | 3.82 | 0.0033 |
|  |  | ASV_1444 | Alphaproteobacteria *Rhizobiales* | 3.82 | 0.0037 |
|  | Unhealthy (n=14) | ASV_119 | **Gammaproteobacteria Ahniella** | 4.24 | 0.0035 |
|  |  | ASV_95 | *Myxococcia Myxococcaceae* | 3.90 | 0.0115 |
|  |  | ASV_40 | **Actinobacteria Mycobacterium** | 3.85 | 0.0250 |

|  |  |  |  |
| --- | --- | --- | --- |
| ASV_334 | Alphaproteobacteria *Sandaracinobacter* | 3.82 | 0.0058 |
| ASV_230 | Gemmatimonadetes *Gemmatimonas* | 3.61 | 0.0014 |

**Table 3.**
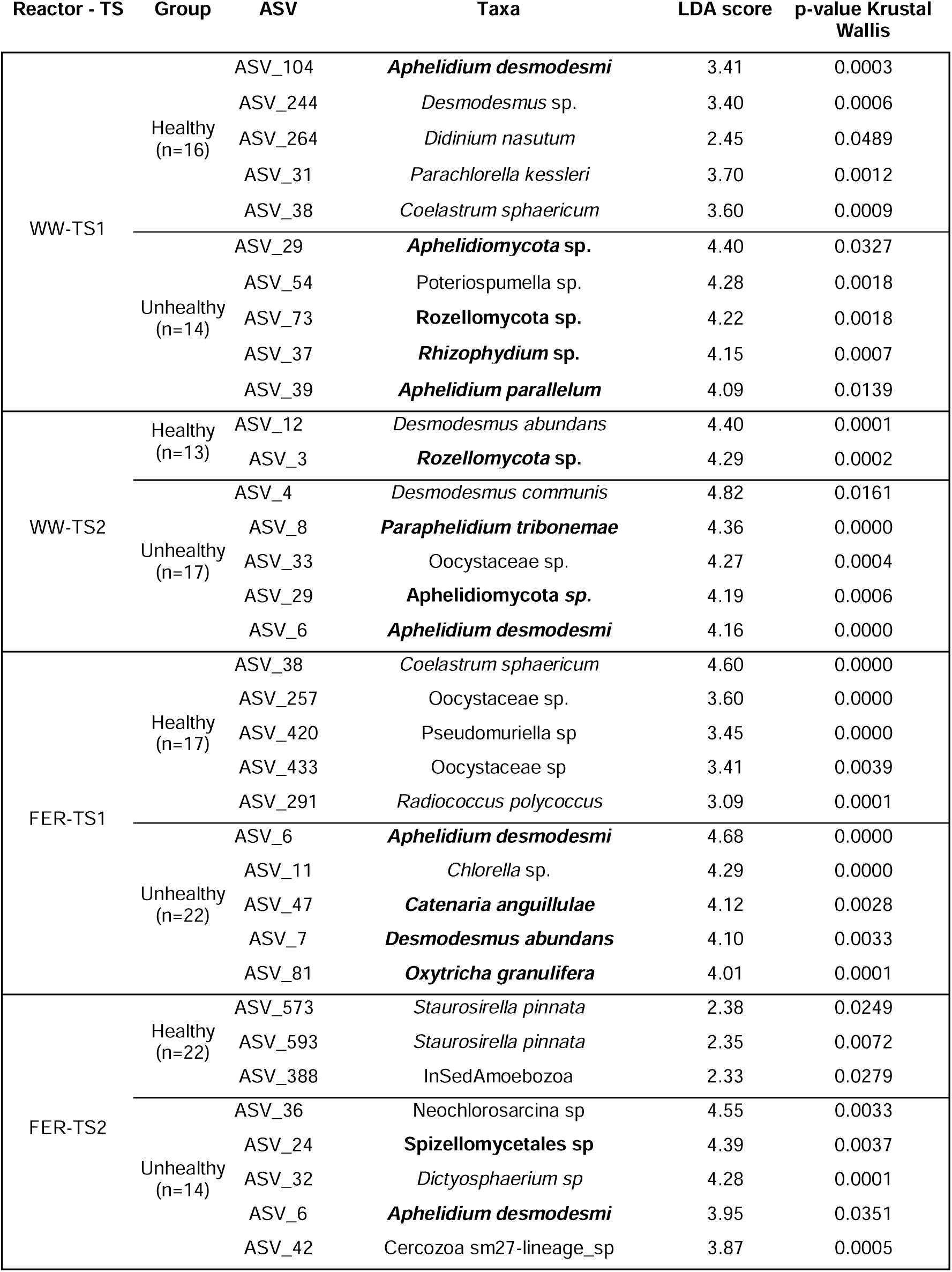
Top-five eukaryotic ASVs related to healthy and unhealthy microbiomes. Healthy (samples in which *Desmodesmus* sp. ’s relative abundance was >70% of the total abundance of eukaryotes) and unhealthy (samples in which Desmodesmus sp. ’s relative abundance was below 20%) microbiome according to LEfSe analysis. Bold text corresponds to the microalgae of study and relevant fungi.

The composition of the eukaryotic communities also varied significantly between reactors (Figures 4 and 5). More than 50% of the eukaryotic ASVs differed between reactors and time series (TS1: X-squared = 8.3386, df = 1, p-value = 0.003881; TS2: X-squared = 7.8368, df = 1, p-value = 0.005119).

The eukaryote community despite being represented by a low percentage of Chlorophyta ASVs (7.6% in see Table 1) presented Chlorophyceae as the most abundant class in both reactors and time series (Table S19). Fungi and other groups dominated the eukaryotic microbial communities in terms of ASV richness (Table 1). As expected, the dominant ASV was *Desmodesmus armatus* (ASV_1), the target microalgae of this study. *D. armatus* was significantly more abundant than other Chlorophyta across all reactors and time series (p-values < 0.001). Other abundant eukaryotic microorganisms included fungi, such as Rozellomycota and Aphelidiomycota, and other groups, such as Cercozoa.

*D. armatus* coexisted with various eukaryotic heterotrophs, including fungal parasites. In our study, the fungal community ranged from 44% of the total eukaryotic abundance reads in the FER reactor to 69% in the WW reactor. The total abundance of fungi was higher in the WW reactor than in the FER reactor, being this difference only statistically significant in TS2 (W = 18796, p-value = 7.091e-06). The most abundant fungi (Figures 4 and 5) included algal endoparasites like Rozellomycota (Letcher et al., 2013) and Aphelidiomycota (Seto et al., 2023), parasites like Chytridiomycota (Laezza et al., 2022) and facultative pathogens like Blastocladiomycota (Asatryan et al., 2019).

The LEfSe analysis also found prevalent eukaryotic ASVs in healthy and unhealthy microbiomes (Table 3). Specifically, fungal ASVs were linked with microbiome health depending on the reactor and time series (Tables 3, S3, and S4).

The Aphelidiomycota *Aphelidium desmodesmi* (ASV_6) appeared always linked with the unhealthy microbiomes of the WW-TS2 and FER-TS1 and TS2. While, another *A. desmodesmi* (ASV_104) was associated with the unhealthy microbiome of the WW-TS2 and FER-TS1, but with the healthy one in WW-TS1 (Tables S5 and S6). Other aphelids, Aphelidiomycota ASV_29 and the Aphelidium parallelum (ASV_39), were linked only with the unhealthy microbiomes of the wastewater reactor during both time series, while the aphelid *Paraphelidium tribonemae* (ASV_8) was linked to the unhealthy microbiome only in WW-TS2. Rozellomycota was represented through different ASVs in the unhealthy microbiome. In FER-TS1 and WW-TS1 we found eight, in FER-TS2 nineteen and in WW-TS2 fourteen (Tables S5 and S6). ASV_3, as the top, was linked to unhealthy microbiomes in WW-TS2 but healthy in FER-TS1. Otherwise, ASV_73 was only found in unhealthy WW-TS1-TS2 and FER-TS1 microbiomes, as well as the ASV_71 being consistently linked with the unhealthy microbiome from FER during both time series. Regarding chytrids, the *Rhizophydium* sp. ASV_37 appeared during the TS1 for both reactors in the unhealthy group. Other ASVs of this genus were also found in the unhealthy microbiome for both reactors and TS. Distinctly, the chytrid Spizellomycetales sp. (ASV_24) only appeared as unhealthy in FER-TS2. Finally, regarding, the Blastocladiomycota class had only *Catenaria anguillulae* ASV_47 linked to unhealthy microbiomes in FER-TS1 and WW-TS2 and ASV_63 only in FER-TS2.

ASVs within the Trebouxiophyceae class, belonging to the Chlorophyta, showed variable associations with the health of the microbiome. While *Coelastrum sphaericum* ASV_38 was linked to the unhealthy in WW-TS2, during FER-TS1 and WW-TS1 was to the healthy. Interestingly, different Oocystaceae ASVs were found in both healthy (ASV_433 and ASV_257 in FER-TS1) and unhealthy (ASV_33 and ASV_257 in WW-TS2) microbiomes. This suggests that Trebouxiophyceae ASVs exhibit a dual behavior, adapting to different conditions and potentially playing different context-dependent roles depending on the overall state of the reactor microbiome.

Ciliates and small flagellates were also detected in both reactors. Members of the phylum Ciliophora, known to be important herbivores in microalgal cultures, were detected at low relative abundances (Table S19). Different Ciliophora ASVs, including *Tokophrya infusorium*, were linked with a healthy microbiome in WW-TS1 but unhealthy in WW-TS2. Yet, *Paramecium polycaryum* (FER-TS1 and TS2) was classified as strongly associated with an unhealthy microbiome (Table S5, S6). No significant decrease in the total number of microalgae and bacteria was observed following their initial appearance in TS1, yet, a potential predator-prey relationship emerged in the WW reactor during TS2. From July to October, the increase in the abundance of ciliates paralleled a decrease in Chlorophyta ASVs, suggesting that ciliates may have been actively feeding on the microalgae.

### Potential microalgae-microbiome interactions

Network analyses revealed similar values for key metrics across the initial, filtered, and *Desmodesmus*-microbiome networks, regardless of reactor or time series (Tables S20-21). Overall, the networks in the WW reactor had more associations than the networks in the FER reactor (see number of associations and number of edges in Tables S20 and S21). Initial and filtered networks exhibited a high complexity; diverse composition (numerous nodes) and highly connected (high average degree). However, the *Desmodesmus*-microbiome networks were very small, with no more than 10 nodes and 9 edges in TS1 and 9 nodes and 8 edges in TS2; and had higher edge density with lower average degree. On the other hand, in all the initial and filtered networks there was a tendency for prokaryotes and eukaryotes to form distinct clusters (positive prokaryote-eukaryote assortativity), meaning prokaryotes tended to connect to each other, and eukaryotes tended to connect to each other. On the contrary, in all the *Desmodesmus*-microbiome networks bacteria and eukaryotes showed a tendency to connect with each other within the network (negative bacterial-eukaryotic assortativity), indicating frequent cross-domain interactions. Lastly, initial and filtered networks exhibited high modularity (values > 0.6), indicating a clustered structure with distinct groups. This suggests the presence of functionally specialized microbial sub-communities within the ecosystem. *Desmodesmus*-microbiome networks had extremely low or almost non-existent modularity, reflecting a poorly defined structure (values close to 0).

We focused our attention on the potential interactions involving *D. armatus* ASV_1, the most abundant algae in the system, and therefore crucial for the microalgae biomass production system (Figure 6). We found up to 10 potential interactions between *D. armatus* and other microbial species in all *Desmodesmus-*microbiome networks (WW and FER, Tables S17 and S18 respectively). Overall, the network analysis revealed a positive association with the Cyanobacteria *Geminocystis* PCC-6308 (ASV_2), and negative associations with the Actinobacteria *Mycobacterium* (ASV_3089, ASV_210, ASV_2037, ASV_34) and with the fungi Rozellomycota (ASV_902) in FER-TS1.

**Figure 6.**
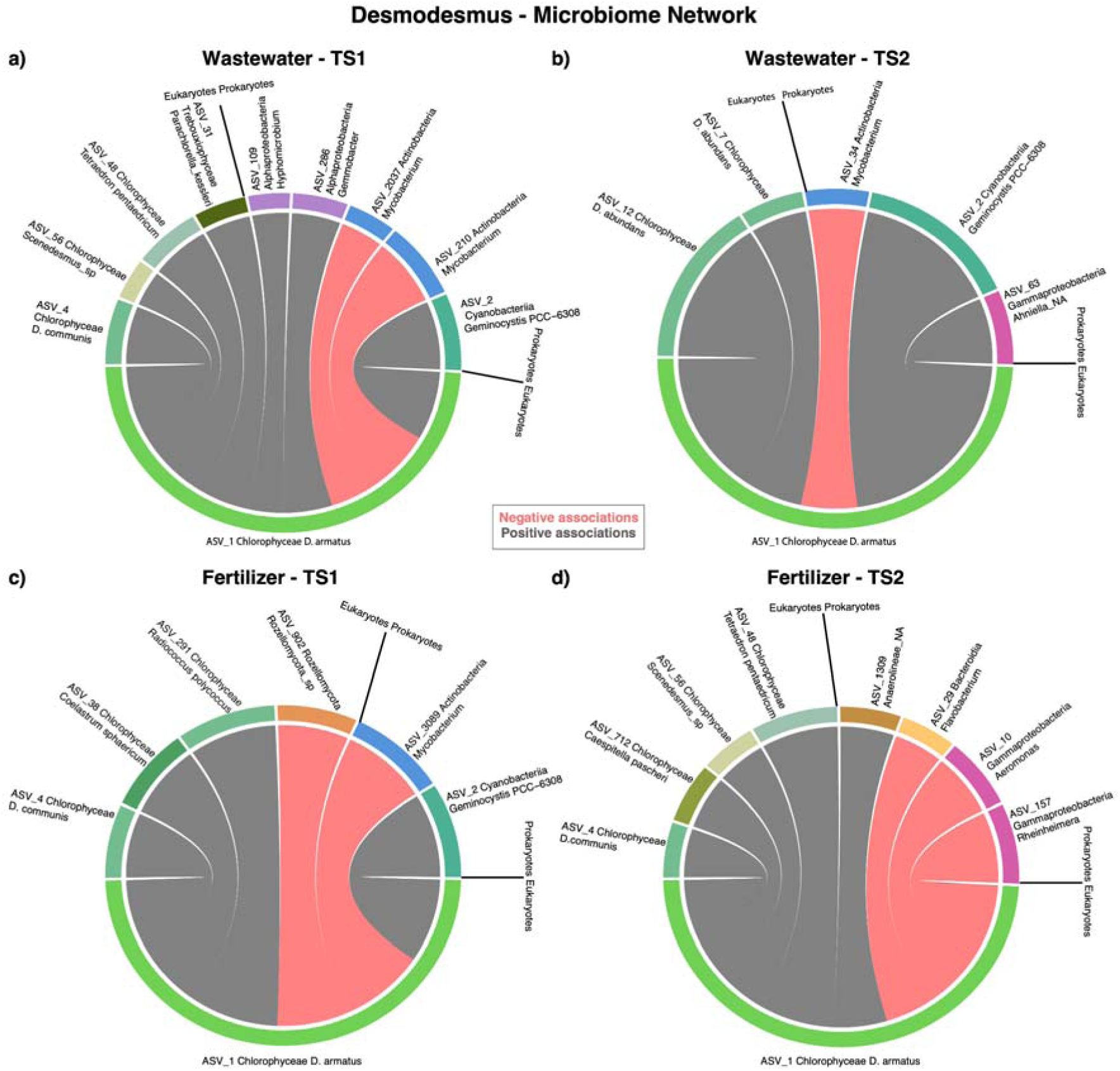
*Desmodesmus*-microbiome networks. Chord diagrams showing the network of interactions/associations between the microalgae and its microbiome in the different reactors and time series at genus level. In these diagrams, the edges (links) show the connection between the ASVs, with the width of the links representing the strength of the interaction. Grey edges are positive associations, while the red are negative ones. The outer colors represent the taxonomic class of the ASVs, and the width of the sectors is indicative of the total number of connections in which a taxonomic class is involved (i.e., the wider the sector, the greater the number of connections in the network).

Regarding the WW reactor (Table S17), we found 7 positive and 2 negative associations in TS1, and 4 positive and 1 negative associations in TS2 (Figure 6a, b). Associations between *D. armatus* and prokaryotes constituted over 60% of all the associations in the WW reactor during both TS. In TS1, there were only two negative associations and were observed between *D. armatus* and the *Mycobacterium* (ASV_210, ASV_2037). In TS2, there was only a negative association, again with the *Mycobacterium* (ASV_34). Among the positive associations identified in the WW reactor (Table S17), the association with the *Geminocystis* PCC-6308 (ASV_2) was the only one prevalent throughout the time series. There was also a positive association with the Gammaproteobacteria *Ahniella* (ASV_63) in WW-TS2, which also appeared representing the healthy microbiome during that period. Other ASVs of these taxa were also linked with the healthy group in WW-TS1 and TS2. *Desmodesmus* (ASV_1) was also positively associated with various ASVs from the classes Trebouxiophyceae and Chlorophyceae. No associations between *D. armatus* and fungi were detected in the WW reactor.

Regarding the FER reactor (Table S18), we found four positive and two negative associations in TS1, and five positive and three negative associations in TS2 (Figure 6c, d). The associations between *D. armatus* and prokaryotes represented approximately 30% in TS1 and 50% in TS2. The positive prokaryotic associations differed between the two-time series (Table S18). During TS1 *D. armatus* exhibited positive associations with the Cyanobacteria *Geminocystis* (ASV_2), but also with eukaryotic organisms such as *Coelastrum*, *Radiococcus* and *D. communis*. (Figure 6c, d). The strongest positive association in FER-TS2 was with *Tetraedron*. Other associations with various ASVs from the Chlorophyceae classes were found. In FER-TS1 reactor, we found only two negative association with the Actinobacteria *Mycobacterium* (ASV_3089) and the fungi Rozellomycota sp. (ASV_902), whereas in TS2, the three negative associations involved the Bacteroidia *Flavobacterium* (ASV_29), the Gammaproteobacteria *Aeromonas* and *Rheinheimera*.

## Discussion

### Richness and dynamics of the microbiome

Our microalgae biomass production raceways harbored highly diverse microbial communities that displayed large variations in species (ASV) richness and community composition over the two 8-month time series (Figures 2 and 3). This is not surprising as open culture systems attain a comparatively higher microbial diversity than closed systems (e.g., thin layer and tubular photobioreactors), in which environmental contamination is minimal and the number of microhabitats is restricted (Bani et al., 2021). Prokaryotic communities were more diverse than their eukaryotic counterparts in both reactors and time series, aligning with previous studies. For example, Bani et al. (2020, 2021) observed that, in *Scenedesmus* cultures under various photobioreactor setups, there were 33% more prokaryotic than eukaryotic ASVs (Bani et al., 2021). Interestingly, the total ASV richness of prokaryotic communities observed in our FER reactors, approximately 2800 ASVs, aligns closely with the values reported by Bani et al. (2020) for systems using tap water. Similarly, the total ASV richness of eukaryotic communities in our FER reactors, approximately 1,000 ASVs, was also consistent with the findings of Bani et al. (2021), who observed a similar richness in clean water systems enriched with fertilizers. Our findings, in conjunction with those from other studies, provide a broad perspective on the range of microbial richness that can be expected in open photobioreactors.

Picoeukaryote and nanoeukaryote communities, particularly in TS2, displayed pronounced compositional turnover, with greater variability in WW reactors and more homogeneity in FER reactors. Prokaryote communities also showed similar trends, with higher heterogeneity in WW reactors. When considering the main differences between WW and FER reactors, the previous results suggest that nutrient availability and the microbial composition of the inflow are likely key drivers of microbial community dynamics within the WW reactors. These patterns align with the notion that nutrient availability impacts microbial community structure and turnover, as highlighted by Ibekwe et al., (2017), Tait et al., (2019) and Ferro et al., (2020). Furthermore, the type of water influences the structure and dynamics of the algae-associated microbiome (Bani et al., 2020; Ferro et al., 2020; Villaró et al., 2022). In turn, FER reactors normally achieve greater stability than WW reactors (Bani et al., 2021).

### Microbiome taxonomic composition

Previous research has demonstrated the significant influence of specific bacterial groups on microalgal growth. For example, Ferro et al. (2020) found that Proteobacteria (specially Alpha and Gamma-proteobacteria), Bacteroidota, Actinobacteriota, and Firmicutes significantly impacted the growth of *Tetradesmus* in wastewater treatment systems. Similarly, Li et al. (2022) reported that Proteobacteria, Bacteroidota, and Cyanobacteria were key prokaryotes influencing the growth of *Chlorella* in a photobioreactor. Our results suggest that *D. armatus* can grow in distinct bacterial communities, whose taxonomic compositions are determined by nutrient regimes and by the taxonomic composition of immigrants carried by the inflowing water. This observation aligns with the findings of Villaró et al. (2022), who found that the composition of the prokaryotic communities in cultures of *Tetradesmus almeriensis* differed among reactors according to the water type used. Our study revealed complex prokaryotic communities in both WW and FER reactors. This complexity likely stems from the influx of microbes through various sources like air, rain, and the influent water itself.

The eukaryotic communities in the reactors were dominated by microalgae and fungi. As expected, the most dominant microorganism was the green microalgae *Desmodesmus armatus*, the culture microalga. Our findings align with those of Carney et al. (2014), who observed a similar eukaryotic community dominated by *Desmodesmus* and *Scenedesmus* (the microalgae from the inoculum) in a mass culture grown in municipal wastewater. This community also included other green algae, fungi, alveolates, and various other taxa.

While Lopes et al., 2023 identified *Chlorella* as a competitor of *Scenedesmus dimorphus*, its low abundance in our study suggests it likely did not compete significantly with *Desmodesmus* for resources. Similarly, although ciliates feed on both microalgae and bacteria and can decimate dense cultures of microalgae in a matter of days (Hadas et al., 1998; Moreno-Garrido and Canavate, 2001; cited in Lakaniemi et al., 2012), their presence did not substantially reduce the abundance of bacteria or microalgae in our time series.

Fungi played a more significant role within the microalgae reactor’s eukaryotic community, particularly during specific periods in the WW treatment. Fungi contribute to the decomposition of organic matter in natural systems, and most likely played a crucial role in carbon and nitrogen cycling within the reactors. However, while fungi can contribute positively to the health of the aquatic ecosystem, they can also act as parasites of microalgae (Laezza et al., 2022; Pan et al., 2023). The parasitic potential of fungi is highlighted by the diversity of fungal groups in microalgal mass culture systems (Table S22). Chytrids are one of the most frequent parasitic fungi associated with microalgae in open systems (Laezza et al., 2022). The diversity and dynamics of fungal communities varied significantly across the reactors in this study. These variations likely arose from a complex interplay of factors, including differences in environmental conditions within each reactor, such as temperature, light intensity, and water flow. Nutrient availability, particularly the types and concentrations of carbon and nitrogen sources, also likely played a role in shaping fungal community composition. Additionally, interactions with other microbes, including bacteria, algae, and other fungi, may have influenced fungal community structure.

### Uncovering microalgae-microbiome interactions

Co-occurrence network analysis revealed predominantly positive interactions between *D. armatus* (ASV_1) and its microbiome in all reactors and TS. Generally, *D. armatus* exhibited slightly more relationships with bacteria than with other eukaryotes. This may have been due to differences in the ASV richness of prokaryotes and eukaryotes, but it is also consistent with the negative bacterial-eukaryotic assortativity values from the *Desmodesmus*-microbiome network, reflecting bacteria’s tendency to connect with eukaryotes, possibly microalgae. Bani et al. (2020) found high interaction values between the most abundant bacterial and eukaryotic OTUs in a microalgae raceway. This is not surprising because microalgae and bacteria have coexisted over evolutionary timescales, mutually interacting to optimize resource use and evolve their life-history traits to varying environmental conditions (Amin et al., 2015; Jiang et al., 2021). Importantly, no fungal associations with *D. armatus* were detected in the WW reactor despite their high relative abundance. This absence suggests that either the interactions with *D. armatus* were not strong enough to be detected, or that, in this specific case, fungi may be interacting more strongly with other eukaryotes or bacteria. These interactions could still indirectly affect *Desmodesmus* in various context-dependent ways, as explained below.

### Identification of potential beneficial bacteria for the microalgae

The present study aimed to identify potential positive and negative interactions between microalgae (*D. armatus)* and its microbiome. Certain bacteria can offer significant benefits (Jiang et al., 2021). It is well known that some cyanobacteria often coexist with microalgae in wastewater treatment reactors (Ibekwe et al., 2017). Our study identified the cyanobacteria *Geminocystis* PCC-6308 (ASV_2) as potentially beneficial to *D. armatus* cultures. This is based on its consistent presence within healthy microbiomes across both reactors and time series, along with its positive associations observed in the microbial networks in both reactors. Potentially beneficial bacteria for *D. armatus* in wastewater reactors might be the Gammaproteobacteria *Ahniella* (ASV_63), which showed a positive interaction with the microalgae and appeared in the healthy microbiome, both in WW-TS2. Moreover, previous studies confirmed that *Ahniella* is a common bacterium in the microalgal biofilms during the co-cultivation of microalgae-activated sludge for municipal wastewater treatment (Cheng et al., 2022) having the capacity to enhance the growth of microalgae in those treatments (Li et al., 2023). The genus *Thiocapsa* (class Gammaproteobacteria) can also be considered as a potentially beneficial microorganism in WW reactors due to its link with the healthy microbiome in TS1 and TS2. *Thiocapsa* is a genus of bacteria that includes purple phototrophic bacteria (Asao et al., 2007). This type of bacteria can benefit *Desmodesmus* by participating in carbon, nitrogen and phosphorus recycling (Hülsen et al., 2018). Another possibly favorable bacterium is *Bosea*, from Rhizobiales class, because of its presence in the healthy microbiome group in FER (TS1 and TS2) and WW (only TS2). The Rhizobiales class include bacteria commonly present in the algal phycosphere. More specifically, Rhizobiales have been observed in bacterial communities of industrial-scale cultures of the microalgae *Tetraselmis suecica* (Piampiano et al., 2019), *Chlorella sorokiniana* (Paddock et al., 2020), *Chlorella vulgaris* (Lakaniemi et al., 2012) and, *Scenedemus dimorphus* (Ferro et al., 2020).

### Identification of potential pathogenic bacteria for the microalgae

The assignment of *Mycobacterium* to an unhealthy microbiome, and its negative co-occurrence with *D. armatus*, underlines its potentially detrimental effect on the growth of *Desmodesmus*. The genus *Mycobacterium* encompasses a wide range of species, including those responsible for tuberculosis and nontuberculous environmental species (Seaman et al., 2022). While pathogenic mycobacteria have been found in marine organisms like fish and mollusks (Carella et al., 2019), their presence in microalgal cultures has not been previously documented. This novel finding underscores the need for further research to understand the role of *Mycobacterium* in microalgal systems and to evaluate their potential impact on the stability and health of these cultures.

*Flavobacterium* (ASV_29) showed a negative co-occurrence with *D. armatus* in FER-TS2. Moreover, its ASV_103 and ASV_240 were consistently linked to an unhealthy microbiome in WW and FER during TS1; while different ASVs were associated with the unhealthy group in both reactors during TS2. However, it was also associated with a healthy microbiome in FER-TS1 and TS2, and WW-TS2, via distinct ASVs. This indicates that *Flavobacterium*’s role in microalgal health may be complex and context-dependent, varying across different systems or conditions. Some *Flavobacterium* organisms have been recognized for decades as a frequent contaminant in large-scale microalgal cultures in open reactors, causing direct adverse effects (Afi et al., 1996). This bacterial genus, which is mostly harmless, can occasionally act as opportunistic or pathogenic (Waśkiewicz & Irzykowska, 2014). In freshwater and marine ecosystems, *Flavobacterium* dominates when organic substrates are present, suggesting that these bacteria may have a specialized role in the uptake, degradation, and decomposition of organic matter (Waśkiewicz & Irzykowska, 2014). Certain flavobacteria species are capable of attacking microalgae through direct cell-to-cell contact or by releasing extracellular compounds that degrade cell walls and destroy DNA (Gómez=Pereira et al., 2012; Laezza et al., 2022; Teeling et al., 2012). Given the evidence, it is reasonable to conclude that *Flavobacterium* ASV_29, ASV_103 and ASV_240 may be detrimental to the microalgae, at least under the specific conditions of this study.

### Identification of microorganisms with diverse roles in the microalgae cultures

Our results highlight the varying roles of fungal groups in *Desmodesmus armatus* cultures. Several fungi, such as Chytridiomycota, Rozellomycota, Aphelidiomycota, and Basidiomycota, were associated with an unhealthy microbiome, indicating their detrimental effects on microalgae. However, some species of these fungi were also found to coexist with *D. armatus* without apparent detrimental effects, suggesting that their roles can shift depending on the biological and environmental conditions (Bittleston et al., 2020).

The Aphelidiomycota *P. tribonemae* (ASV_8), identified as an indicator of an unhealthy microbiome, proliferated during periods of low microalgal abundance in WW-TS2. This suggests a potential pathogenic role for *P. tribonemae* in *D. armatus* cultures. While this aphelid was first described as a parasite of the alga *Tribonema gayanum* (Karpov et al.), it has never been reported to infect *D. armatus*. This finding may represent the first evidence of *P. tribonemae* as a pathogen for *D. armatus*. The aphelid Aphelidiomycota sp. (ASV_29) and *Aphelidium parallelum* (ASV_39) were constantly linked to the unhealthy microbiome of the WW reactor for both TS. Seto et al., (2022) found that *A. parallelum* parasitizes *Monoraphidium* sp. and other strains of Selenastraceae. There is no literature on *A. parallelum* parasitizing *D. armatus*, so this study may represent the first evidence of its detrimental effect on *D. armatus*. On the other hand, *Aphelidium desmodesmi* (ASV_104 and ASV_6), linked to the unhealthy microbiome in FER-TS1-TS2 and WW-TS2, is a known parasite of *D. armatus*. Yet, *A. desmodesmi* (ASV_104) was also linked to the healthy microbiome in WW-TS1 suggesting a more complex relationship. Letcher and colleagues (2017) reported this fungal pathogen, along with *Aphelidium chlorococcarum*, infecting *D. armatus* cultures (Letcher et al., 2013, 2015). Thus, only the *A. desmodesmi* (ASV_6) would be considered as likely damaging to the microalgae. These results underscore the context-dependent role of *A. desmodesmi* in the microalgal community, where its impact can vary depending on the specific environment and interacting organisms present (Bittleston et al., 2020). These findings align with previous studies showing that fungi can act as both competitors and symbionts in microalgae cultures (Laezza et al., 2022). Similarly, Rozellomycota sp. (ASV_3) also exhibited contrasting dynamics, being linked with the unhealthy microbiome and having a negative association with the microalgae in FER-TS1 but with the healthy microbiome in WW-TS2. While ASV_71 and ASV_73 were consistent in unfavorables microbiomes throughout reactors and TS. This is expected, as Rozellomycota, previously known as Cryptomycota, are parasites of microalgae (Karpov et al., 2017; Torruella et al., 2018). Previous reports have shown species of Rozellomycota limiting the growth of *Scenedesmus* sp. in both laboratory reactors and open raceways used for mass cultivation (Gromov & Mamkaeva, 1970; P. M. Letcher et al., 2013). The presence of the chytrid *Rhizophydium* sp. (ASV_37) in the unhealthy microbiome of WW-FER-TS1 further supports the detrimental role this genus plays in microalgal health (Letcher & Powell, 2012). Similar results have been reported for the species *Rhizophydium scenedesmi* (Ding et al., 2018). *R. chlorogonii* and *Rhizophydium* sp. have already been identified as dominant fungi in photobioreactors with *Desmodesmus* and *Scenedesmus* (Carney et al., 2014). Similarly, Spizellomycetales (ASV_24), at the top fungi in the unhealthy group of FER-TS2, it’s a chytrid that include saprophytes and/or parasites that have been found in freshwater and soil environments as in antarctic lake sediments (Gonçalves et al., 2022) but there’s no literature about its presence in microalgae cultures or affecting microalgae in natural environments. Therefore, this is not sufficient evidence to conclude that it could be harmful to the microalga. Finally, the Blastocladiomycota *Catenaria anguillulae*, found in the unhealthy microbiome in FER-TS1-TS2 and WW-TS2, is known for being a parasite for plants (Gleason et al., 2014) but not for microalgae. So, we cannot determine its harmful effect on microalgae cultures.

Understanding the intricate interplay between fungi and microalgae is crucial for optimizing microalgal cultivation and harnessing the full potential of these systems. Fungal parasitic interactions occur during specific stages of their life cycle when they attach, encyst, germinate, and produce rhizoids that penetrate their hosts and release digestive enzymes, completing their parasitic cycle (Lauritano & Galasso, 2023). However, under certain conditions, they may exhibit a neutral or even positive influence on microalgae. This can be attributed to factors such as (1) their presence as zoospores, a motile stage where they swim freely in search of host cells without causing harm before attachment (Laezza et al., 2022; Lauritano & Galasso, 2023); (2) specific biological and environmental factors that can potentially modulate microbial interactions within the community (Bittleston et al., 2020); (3) indirect effects, where fungi may parasite other eukaryotes that are competitors of the culture algae, or bacteria detrimental to *D.armatus* but also (4) where fungi contribute to decompose organic matter, thereby releasing minerals and nutrients essential for the microalgae (Wang et al., 2022; Lauritano & Galasso, 2023). These observations underscore the dynamic and context-dependent nature of fungal interactions within microalgal cultures. In summary, fungal communities can shift between beneficial, neutral, and detrimental roles depending on a variety of factors, including the specific fungal species present, the microalgae, and the prevailing environmental conditions. This variability highlights the complexity of these interactions and the need for a deeper understanding of the ecological factors that govern fungal dynamics in microalgal systems.

## Conclusions

Large-scale microalgal raceway reactors host a remarkably diverse microbiome. Our analysis unveiled the dynamic and complex nature of this microbiota, characterized by intricate interactions between microalgae and their associated microorganisms, with significant implications for microalgal growth and productivity. Co-occurrence network analysis revealed a complex network of interconnections within the microbial community. Interactions between eukaryotic and bacterial communities offered valuable insights into the ecological dynamics of these reactor systems and highlighted a critical knowledge gap in our understanding of their interactome. Our study revealed a diverse bacterial community dominated by Alpha- and Gammaproteobacteria, Bacteroidia, Cyanobacteria, and Actinobacteria. Within this community, *Geminocystis*, *Thiocapsa, Ahniella* and *Bosea* were associated with a healthy microbiome, potentially promoting the growth of the target microalga, *Desmodesmus armatus*. Conversely, known pathogens like *Mycobacterium* sp. and *Flavobacterium* were linked to an unhealthy microbiome. Among eukaryotes, fungi exhibited dynamic and context-dependent associations. However, some showed consistent unfavorable dynamics, with *Paraphelidium tribonemae* (ASV_8) and *Aphelidium parallelum* (ASV_39) being identified for the first time as potential parasites of *D. armatus*, while *A. desmodesmi* (ASV_6), Aphelidiomycota sp. (ASV_29), Rozellomycota sp. (ASV_73 and ASV_71) and *Rhizophidium* sp. (ASV_37) being reaffirmed as harmful organisms. Further research is crucial to confirm these interactions and assess their broader applicability across diverse microalgal cultivation systems. Ultimately, high-frequency monitoring of microbiome composition and dynamics is essential to understanding these complex interactions, their role in cultivation systems, and potential applications for enhancing microalgal biomass production.

## CRediT authorship contribution statement

**Judith Traver-Azuara**: Conceptualization. Investigation: sample treatment. Data Curation: produce, scrub and maintain research data, produce metadata. Formal analysis: statistical and computational analyses to the data. Software: designing and implementation of the computer code. Methodology. Visualization. Writing – original draft. **Caterina R. Giner:** Resources. Data curation. Formal analysis. **Carmen Garcia-Comas:** Conceptualization. Investigation. Supervision: supervision of statistical and computational analyses. Writing – review & editing. **Ana Sánchez-Zurano, Martina Ciardi, and Gabriel Acién:** Resources. Investigation: perform experiments and data collection. Data Curation: produce metadata. **Aleix Obiol and Ramon Massana:** Data Curation: scrub data, eukaryotic taxonomy. Writing – review & editing: revision. **Sofiya Bondarenko:** Data Curation: fungi database curation. Writing – review & editing: revision. **Montse Sala:** Writing – review & editing: revision. **R.Logares:** Resources. Conceptualization. Investigation. Supervision. Data curation. Software: bioinformatics and interaction networks. Writing – review & editing. **P. Cermeno:** Conceptualization. Investigation. Funding acquisition. Supervision. Writing – review & editing.

All authors have read and agreed to the published version of the manuscript.

## Acknowledgments

We thank the Marine Bioinformatics Service (MARBITS; http://marbits.icm.csic.es) of the Institut de Ciències del Mar—CSIC and the Drago supercomputer of the CSIC for providing the computing power needed to perform some of the bioinformatic analyses. We thank the team running the microalgae culturing facility at the University of Almeria.

This work has been supported by the European Union’s Horizon 2020 Research and Innovation Programme through the project PRODIGIO (N. 101007006): “Developing early-warning systems for improved microalgae PROduction and anaerobic DIGestION”, and by the project INCEPTION “In-depth characterization of microalgal crop genomics for sustainable biomass production and circular bioeconomy” (TED2021-131071B-I00), financed by the Spanish Research Agency (AEI). This work also acknowledges the Severo Ochoa Centre of Excellence accreditation (CEX2019-000928-S).

## Data availability

Data will be released upon acceptance.

## References

Afi, L., Metzger, P., Largeau, C., Connan, J., Berkaloff, C., & Rousseau, B. (1996). Bacterial degradation of green microalgae: Incubation of Chlorella emersonii and Chlorella vulgaris with Pseudomonas oleovorans and Flavobacterium aquatile. 25, 117–130.

Ahmad, A., & Ashraf, S. S. (2023). Sustainable food and feed sources from microalgae: Food security and the circular bioeconomy. Algal Research, 74, 103185. 10.1016/j.algal.2023.103185

Alam, Md. A., Wan, C., Tran, D. T., Mofijur, M., Ahmed, S. F., Mehmood, M. A., Shaik, F., Vo, D.-V. N., & Xu, J. (2022). Microalgae binary culture for higher biomass production, nutrients recycling, and efficient harvesting: A review. Environmental Chemistry Letters, 20(2), 1153–1168. 10.1007/s10311-021-01363-z

Amin, S. A., Hmelo, L. R., Van Tol, H. M., Durham, B. P., Carlson, L. T., Heal, K. R., Morales, R. L., Berthiaume, C. T., Parker, M. S., Djunaedi, B., Ingalls, A. E., Parsek, M. R., Moran, M. A., & Armbrust, E. V. (2015). Interaction and signalling between a cosmopolitan phytoplankton and associated bacteria. Nature, 522(7554), 98–101. 10.1038/nature14488

Asao, M., Takaichi, S., & Madigan, M. T. (2007). Thiocapsa imhoffii, sp. Nov., an alkaliphilic purple sulfur bacterium of the family Chromatiaceae from Soap Lake, Washington (USA). Archives of Microbiology, 188(6), 665–675. 10.1007/s00203-007-0287-9

Asatryan, A., Boussiba, S., & Zarka, A. (2019). Stimulation and isolation of Paraphysoderma sedebokerense (Blastocladiomycota) propagules and their infection capacity toward their host under different physiological and environmental conditions. Frontiers in Cellular and Infection Microbiology, 9, 72. 10.3389/fcimb.2019.00072

Auguie, B., & Antonov, A. (2017). gridExtra v.2.3. 10.32614/CRAN.package.gridExtra

Awasthi, A., Singh, M., Soni, S. K., Singh, R., & Kalra, A. (2014). Biodiversity acts as insurance of productivity of bacterial communities under abiotic perturbations. The ISME Journal, 8(12), 2445–2452. 10.1038/ismej.2014.91

Balzano, S., Abs, E., & Leterme, S. (2015). Protist diversity along a salinity gradient in a coastal lagoon. Aquatic Microbial Ecology, 74(3), 263–277. 10.3354/ame01740

Bani, A., Fernandez, F. G. A., D’Imporzano, G., Parati, K., & Adani, F. (2021). Influence of photobioreactor set-up on the survival of microalgae inoculum. Bioresource Technology, 320, 124408. 10.1016/j.biortech.2020.124408

Bani, A., Parati, K., Pozzi, A., Previtali, C., Bongioni, G., Pizzera, A., Ficara, E., & Bellucci, M. (2020). Comparison of the performance and microbial community structure of two outdoor pilot-scale photobioreactors treating digestate. Microorganisms, 8(11), 1754. 10.3390/microorganisms8111754

Berg, G., Rybakova, D., Fischer, D., Cernava, T., Vergès, M.-C. C., Charles, T., Chen, X., Cocolin, L., Eversole, K., Corral, G. H., Kazou, M., Kinkel, L., Lange, L., Lima, N., Loy, A., Macklin, J. A., Maguin, E., Mauchline, T., McClure, R., … Schloter, M. (2020). Microbiome definition re-visited: Old concepts and new challenges. Microbiome, 8(1), 103. 10.1186/s40168-020-00875-0

Bittleston, L. S., Gralka, M., Leventhal, G. E., Mizrahi, I., & Cordero, O. X. (2020). Context-dependent dynamics lead to the assembly of functionally distinct microbial communities. Nature Communications, 11(1), 1440. 10.1038/s41467-020-15169-0

Borowitzka, M. A., & Vonshak, A. (2017). Scaling up microalgal cultures to commercial scale. European Journal of Phycology, 52(4), 407–418. 10.1080/09670262.2017.1365177

Callahan, B. J., McMurdie, P. J., Rosen, M. J., Han, A. W., Johnson, A. J. A., & Holmes, S. P. (2016). DADA2: High-resolution sample inference from Illumina amplicon data. Nature Methods, 13(7), 581–583. 10.1038/nmeth.3869

Carella, F., Aceto, S., Pollaro, F., Miccio, A., Iaria, C., Carrasco, N., Prado, P., & De Vico, G. (2019). A mycobacterial disease is associated with the silent mass mortality of the pen shell Pinna nobilis along the Tyrrhenian coastline of Italy. Scientific Reports, 9(1), 2725. 10.1038/s41598-018-37217-y

Carney, L. T., Reinsch, S. S., Lane, P. D., Solberg, O. D., Jansen, L. S., Williams, K. P., Trent, J. D., & Lane, T. W. (2014). Microbiome analysis of a microalgal mass culture growing in municipal wastewater in a prototype OMEGA photobioreactor. Algal Research, 4, 52–61. 10.1016/j.algal.2013.11.006

Cheng, Y., Wang, H., Deng, Z., Wang, J., Liu, Z., Chen, Y., Ma, Y., Li, B., Yang, L., Zhang, Z., & Wu, L. (2022). Efficient removal of Imidacloprid and nutrients by algae-bacteria biofilm reactor (ABBR) in municipal wastewater: Performance, mechanisms and the importance of illumination. Chemosphere, 305, 135418. 10.1016/j.chemosphere.2022.135418

Chhandama, M. V. L., Satyan, K. B., Changmai, B., Vanlalveni, C., & Rokhum, S. L. (2021). Microalgae as a feedstock for the production of biodiesel: A review. Bioresource Technology Reports, 15, 100771. 10.1016/j.biteb.2021.100771

Csárdi, G., Nepusz, T., Traag, V., Horvát, S., Zanini, F., Noom, D., & Müller, K. (2023). Igraph v.1.4.3. 10.32614/CRAN.package.igraph

D’Alessandro, E. B., & Antoniosi Filho, N. R. (2016). Concepts and studies on lipid and pigments of microalgae: A review. Renewable and Sustainable Energy Reviews, 58, 832–841. 10.1016/j.rser.2015.12.162

Das, P. K., Rani, J., Rawat, S., & Kumar, S. (2022). Microalgal co-cultivation for biofuel production and bioremediation: current status and benefits. BioEnergy Research, 15(1), 1–26. 10.1007/s12155-021-10254-8

Day, J. G., Gong, Y., & Hu, Q. (2017). Microzooplanktonic grazers – A potentially devastating threat to the commercial success of microalgal mass culture. Algal Research, 27, 356–365. 10.1016/j.algal.2017.08.024

Delgado-Baquerizo, M., Maestre, F. T., Reich, P. B., Jeffries, T. C., Gaitan, J. J., Encinar, D., Berdugo, M., Campbell, C. D., & Singh, B. K. (2016). Microbial diversity drives multifunctionality in terrestrial ecosystems. Nature Communications, 7(1), 10541. 10.1038/ncomms10541

Deutschmann, I. M., Delage, E., Giner, C. R., Sebastián, M., Poulain, J., Arístegui, J., Duarte, C. M., Acinas, S. G., Massana, R., Gasol, J. M., Eveillard, D., Chaffron, S., & Logares, R. (2024). Disentangling microbial networks across pelagic zones in the tropical and subtropical global ocean. Nature Communications, 15(1), 126. 10.1038/s41467-023-44550-y

Ding, Y., Peng, X., Wang, Z., Wen, X., Geng, Y., Zhang, D., & Li, Y. (2018). Occurrence and characterization of an epibiotic parasite in cultures of oleaginous microalga Graesiella sp. WBG-1. Journal of Applied Phycology, 30(2), 819–830. 10.1007/s10811-017-1302-4

Dolganyuk, V., Belova, D., Babich, O., Prosekov, A., Ivanova, S., Katserov, D., Patyukov, N., & Sukhikh, S. (2020). Microalgae: a promising source of valuable bioproducts. Biomolecules, 10(8), 1153. 10.3390/biom10081153

Ferro, L., Hu, Y. O. O., Gentili, F. G., Andersson, A. F., & Funk, C. (2020). DNA metabarcoding reveals microbial community dynamics in a microalgae-based municipal wastewater treatment open photobioreactor. Algal Research, 51, 102043. 10.1016/j.algal.2020.102043

Fierer, N. (2017). Embracing the unknown: Disentangling the complexities of the soil microbiome. Nature Reviews Microbiology, 15(10), 579–590. 10.1038/nrmicro.2017.87

Gleason, F. H., Lilje, O., Marano, A. V., Sime-Ngando, T., Sullivan, B. K., Kirchmair, M., & Neuhauser, S. (2014). Ecological functions of zoosporic hyperparasites. Frontiers in Microbiology, 5. 10.3389/fmicb.2014.00244

Gómez-Pereira, P. R., Schüler, M., Fuchs, B. M., Bennke, C., Teeling, H., Waldmann, J., Richter, M., Barbe, V., Bataille, E., Glöckner, F. O., & Amann, R. (2012). Genomic content of uncultured Bacteroidetes from contrasting oceanic provinces in the North Atlantic Ocean. Environmental Microbiology, 14(1), 52–66. 10.1111/j.1462-2920.2011.02555.x

Gonçalves, V. N., De Souza, L. M. D., Lirio, J. M., Coria, S. H., Lopes, F. A. C., Convey, P., Carvalho-Silva, M., De Oliveira, F. S., Câmara, P. E. A. S., & Rosa, L. H. (2022). Diversity and ecology of fungal assemblages present in lake sediments at Clearwater Mesa, James Ross Island, Antarctica, assessed using metabarcoding of environmental DNA. Fungal Biology, 126(10), 640–647. 10.1016/j.funbio.2022.08.002

Gouy M, Tannier E, Comte N & Parsons DP. (2021). Seaview Version 5: A Multiplatform Software for Multiple Sequence Alignment, Molecular Phylogenetic Analyses, and Tree Reconciliation. Multiple Sequence Alignment: Methods and Protocols, 2231, 241–260. Springer US. 10.1007/978-1-0716-1036-7

Gromov, B., & Mamkaeva, K. (1970). The fine structure of Amoeboaphelidium protococcarum Gromov et Mamkaeva–an endoparasite of green alga Scenedesmus. 67, 452–459.

Grube, M., Cernava, T., Soh, J., Fuchs, S., Aschenbrenner, I., Lassek, C., Wegner, U., Becher, D., Riedel, K., Sensen, C. W., & Berg, G. (2015). Exploring functional contexts of symbiotic sustain within lichen-associated bacteria by comparative omics. The ISME Journal, 9(2), 412–424. 10.1038/ismej.2014.138

Gu, Z. (2022). Circlize v.0.15. 10.32614/CRAN.package.circlize

Guillou, L., Bachar, D., Audic, S., Bass, D., Berney, C., Bittner, L., Boutte, C., Burgaud, G., De Vargas, C., Decelle, J., Del Campo, J., Dolan, J. R., Dunthorn, M., Edvardsen, B., Holzmann, M., Kooistra, W. H. C. F., Lara, E., Le Bescot, N., Logares, R., … Christen, R. (2012). The Protist Ribosomal Reference database (PR2): A catalog of unicellular eukaryote Small Sub-Unit rRNA sequences with curated taxonomy. Nucleic Acids Research, 41(D1), D597–D604. 10.1093/nar/gks1160

Hülsen, T., Hsieh, K., Lu, Y., Tait, S., & Batstone, D. J. (2018). Simultaneous treatment and single cell protein production from agri-industrial wastewaters using purple phototrophic bacteria or microalgae – A comparison. Bioresource Technology, 254, 214–223. 10.1016/j.biortech.2018.01.032

Ibekwe, A. M., Murinda, S. E., Murry, M. A., Schwartz, G., & Lundquist, T. (2017). Microbial community structures in high-rate algae ponds for bioconversion of agricultural wastes from livestock industry for feed production. Science of The Total Environment, 580, 1185–1196. 10.1016/j.scitotenv.2016.12.076

Jiang, L., Li, Y., & Pei, H. (2021). Algal–bacterial consortia for bioproduct generation and wastewater treatment. Renewable and Sustainable Energy Reviews, 149, 111395. 10.1016/j.rser.2021.111395

Karpov, S. A., Tcvetkova, V. S., Mamkaeva, M. A., Torruella, G., Timpano, H., Moreira, D., Mamanazarova, K. S., & López-García, P. (2017). Morphological and genetic diversity of Opisthosporidia: new aphelid Paraphelidium tribonemae gen. et sp. nov. Journal of Eukaryotic Microbiology, 64(2), 204–212. 10.1111/jeu.12352

Khan, M. I., Shin, J. H., & Kim, J. D. (2018). The promising future of microalgae: Current status, challenges, and optimization of a sustainable and renewable industry for biofuels, feed, and other products. Microbial Cell Factories, 17(1), 36. 10.1186/s12934-018-0879-x

Kim Hue, N. T., Deruyck, B., Decaestecker, E., Vandamme, D., & Muylaert, K. (2018). Natural chemicals produced by marine microalgae as predator deterrents can be used to control ciliates contamination in microalgal cultures. Algal Research, 29, 297–303. 10.1016/j.algal.2017.11.036

Kusmayadi, A., Leong, Y. K., Yen, H.-W., Huang, C.-Y., & Chang, J.-S. (2021). Microalgae as sustainable food and feed sources for animals and humans – Biotechnological and environmental aspects. Chemosphere, 271, 129800. 10.1016/j.chemosphere.2021.129800

Laezza, C., Salbitani, G., & Carfagna, S. (2022). Fungal contamination in microalgal cultivation: biological and biotechnological aspects of fungi-microalgae interaction. Journal of Fungi, 8(10), 1099. 10.3390/jof8101099

Lakaniemi, A.-M., Hulatt, C. J., Wakeman, K. D., Thomas, D. N., & Puhakka, J. A. (2012). Eukaryotic and prokaryotic microbial communities during microalgal biomass production. Bioresource Technology, 124, 387–393. 10.1016/j.biortech.2012.08.048

Lauritano, C., & Galasso, C. (2023). Microbial interactions between marine microalgae and fungi: from chemical ecology to biotechnological possible applications. Marine Drugs, 21(5), 310. 10.3390/md21050310

Letcher, P. M., Lopez, S., Schmieder, R., Lee, P. A., Behnke, C., Powell, M. J., & McBride, R. C. (2013). Characterization of Amoeboaphelidium protococcarum, an algal parasite new to the Cryptomycota isolated from an outdoor algal pond used for the production of biofuel. PLoS ONE, 8(2), e56232. 10.1371/journal.pone.0056232

Letcher, P. M., Powell, M. J., & Davis, W. J. (2015). A new family and four new genera in Rhizophydiales (Chytridiomycota). Mycologia, 107(4), 808–830. 10.3852/14-280

Letcher, P. M., Powell, M. J., Lee, P. A., Lopez, S., & Burnett, M. (2017). Molecular phylogeny and ultrastructure of Aphelidium desmodesmi, a new species in Aphelida (Opisthosporidia). Journal of Eukaryotic Microbiology, 64(5), 655–667. 10.1111/jeu.12401

Letcher, P. M.& Powell, M. J. (2012). A taxonomic summary and revision of Rhizophydium (Rhizophydiales, Chytridiomycota). Tuscaloosa, AL, USA: University Printing, The University of Alabama

Li, S., Chu, Y., Xie, P., Xie, Y., Chang, H., & Ho, S.-H. (2022). Insights into the microalgae-bacteria consortia treating swine wastewater: Symbiotic mechanism and resistance genes analysis. Bioresource Technology, 349, 126892. 10.1016/j.biortech.2022.126892

Li, X., Liu, J., Tian, J., Pan, Z., Chen, Y., Ming, F., Wang, R., Wang, L., Zhou, H., Li, J., & Tan, Z. (2023). Co-cultivation of microalgae-activated sludge for municipal wastewater treatment: Exploring the performance, microbial co-occurrence patterns, microbiota dynamics and function during the startup stage. Bioresource Technology, 374, 128733. 10.1016/j.biortech.2023.128733

Lian, J., Wijffels, R. H., Smidt, H., & Sipkema, D. (2018). The effect of the algal microbiome on industrial production of microalgae. Microbial Biotechnology, 11(5), 806–818. 10.1111/1751-7915.13296

Lopes, G. B., Goelzer, A., Reichel, T., De Resende, M. L. V., & Duarte, W. F. (2023). Potential of Desmodesmus abundans as biofertilizer in common bean (Phaseolus vulgaris L.). Biocatalysis and Agricultural Biotechnology, 49, 102657. 10.1016/j.bcab.2023.102657

Martin, M. (2011). Cutadapt removes adapter sequences from high-throughput sequencing reads. 17, 10–12. 10.14806/ej.17.1.200

Molina-Grima, E., García-Camacho, F., Acién-Fernández, F. G., Sánchez-Mirón, A., Plouviez, M., Shene, C., & Chisti, Y. (2022). Pathogens and predators impacting commercial production of microalgae and cyanobacteria. Biotechnology Advances, 55, 107884. 10.1016/j.biotechadv.2021.107884

Muñoz, R., Guieysse, B., & Mattiasson, B. (2003). Phenanthrene biodegradation by an algal-bacterial consortium in two-phase partitioning bioreactors. Applied Microbiology and Biotechnology, 61(3), 261–267. 10.1007/s00253-003-1231-9

Novoveská, L., Nielsen, S. L., Eroldoğan, O. T., Haznedaroglu, B. Z., Rinkevich, B., Fazi, S., Robbens, J., Vasquez, M., & Einarsson, H. (2023). Overview and challenges of large-scale cultivation of photosynthetic microalgae and cyanobacteria. Marine Drugs, 21(8), 445. 10.3390/md21080445

Obiol, A., Giner, C. R., Sánchez, P., Duarte, C. M., Acinas, S. G., & Massana, R. (2020). A metagenomic assessment of microbial eukaryotic diversity in the global ocean. Molecular Ecology Resources, 20(3), 718–731. 10.1111/1755-0998.13147

Oksanen, J., F. Guillaume, B., Friendly, M., Kindt, R., Legendre, P., McGlinn, D., et al. (2020). vegan: Community Ecology Package. R package version 2.5–7. https://CRAN.R-project.org/package=vegan

Paddock, M. B., Fernández-Bayo, J. D., & Vander Gheynst, J. S. (2020). The effect of the microalgae-bacteria microbiome on wastewater treatment and biomass production. Applied Microbiology and Biotechnology, 104(2), 893–905. 10.1007/s00253-019-10246-x

Pan, X., Yue, Z., She, Z., He, X., Wang, S., Chuai, X., & Wang, J. (2023). Eukaryotic community structure and interspecific interactions in a stratified acidic pit lake water in Anhui province. Microorganisms, 11(4), 979. 10.3390/microorganisms11040979

Parada, A. E., Needham, D. M., & Fuhrman, J. A. (2016). Every base matters: Assessing small subunit rRNA primers for marine microbiomes with mock communities, time series and global field samples. Environmental Microbiology, 18(5), 1403–1414. 10.1111/1462-2920.13023

Piampiano, E., Pini, F., Biondi, N., Pastorelli, R., Giovannetti, L., & Viti, C. (2019). Analysis of microbiota in cultures of the green microalga Tetraselmis suecica. European Journal of Phycology, 54(3), 497–508. 10.1080/09670262.2019.1606940

Quast, C., Pruesse, E., Yilmaz, P., Gerken, J., Schweer, T., Yarza, P., Peplies, J., & Glöckner, F. O. (2012). The SILVA ribosomal RNA gene database project: Improved data processing and web-based tools. Nucleic Acids Research, 41(D1), D590–D596. 10.1093/nar/gks1219

R Core Team (2022). R: A language and environment for statistical computing., ed. R. F. for S. Computing Vienna, Austria.

Rstudio Team (2022). RStudio: Integrated Development Environment for R., ed. P. RStudio Boston, MA.

Ramanan, R., Kim, B.-H., Cho, D.-H., Oh, H.-M., & Kim, H.-S. (2016). Algae–bacteria interactions: Evolution, ecology and emerging applications. Biotechnology Advances, 34(1), 14–29. 10.1016/j.biotechadv.2015.12.003

Rodrigues Reis, C. E., Ogero D’Otaviano, L., Rajendran, A., & Hu, B. (2018). Co-culture of filamentous feed-grade fungi and microalgae as an alternative to increase feeding value of ethanol coproducts. Fermentation, 4(4), 86. 10.3390/fermentation4040086

Schloss, P. D., Westcott, S. L., Ryabin, T., Hall, J. R., Hartmann, M., Hollister, E. B., Lesniewski, R. A., Oakley, B. B., Parks, D. H., Robinson, C. J., Sahl, J. W., Stres, B., Thallinger, G. G., Van Horn, D. J., & Weber, C. F. (2009). Introducing mothur: open-source, platform-independent, community-supported software for describing and comparing microbial communities. Applied and Environmental Microbiology, 75(23), 7537–7541. 10.1128/AEM.01541-09

Seaman, J. L., Oosthuizen, C. B., Gibango, L., & Lall, N. (2022). Mycobacterial quorum quenching and biofilm inhibition potential of medicinal plants. In Medicinal Plants as Anti-Infectives (pp. 309–333). Elsevier. 10.1016/B978-0-323-90999-0.00008-2

Segata, N., Izard, J., Waldron, L., Gevers, D., Miropolsky, L., Garrett, W. S., & Huttenhower, C. (2011). Metagenomic biomarker discovery and explanation. 10.1186/gb-2011-12-6-r60.

Seto, K., Simmons, D. R., Quandt, C. A., Frenken, T., Dirks, A. C., Clemons, R. A., McKindles, K. M., McKay, R. M. L., & James, T. Y. (2023). A combined microscopy and single-cell sequencing approach reveals the ecology, morphology, and phylogeny of uncultured lineages of zoosporic fungi. *mBio*, e01313–23. 10.1128/mbio.01313-23

Seymour, J. R., Amin, S. A., Raina, J.-B., & Stocker, R. (2017). Zooming in on the phycosphere: The ecological interface for phytoplankton–bacteria relationships. Nature Microbiology, 2(7), 17065. 10.1038/nmicrobiol.2017.65

Smith, V. H. (2007). Microbial diversity–productivity relationships in aquatic ecosystems: Diversity–productivity relationships. FEMS Microbiology Ecology, 62(2), 181–186. 10.1111/j.1574-6941.2007.00381.x

Subashchandrabose, S. R., Ramakrishnan, B., Megharaj, M., Venkateswarlu, K., & Naidu, R. (2011). Consortia of cyanobacteria/microalgae and bacteria: Biotechnological potential. Biotechnology Advances, 29(6), 896–907. 10.1016/j.biotechadv.2011.07.009

Tackmann, J., Matias Rodrigues, J. F., & Von Mering, C. (2019). Rapid inference of direct interactions in large-scale ecological networks from heterogeneous microbial sequencing data. Cell Systems, 9(3), 286–296.e8. 10.1016/j.cels.2019.08.002

Tait, K., White, D. A., Kimmance, S. A., Tarran, G., Rooks, P., Jones, M., & Llewellyn, C. A. (2019). Characterisation of bacteria from the cultures of a Chlorella strain isolated from textile wastewater and their growth enhancing effects on the axenic cultures of Chlorella vulgaris in low nutrient media. Algal Research, 44, 101666. 10.1016/j.algal.2019.101666

Teeling, H., Fuchs, B. M., Becher, D., Klockow, C., Gardebrecht, A., Bennke, C. M., Kassabgy, M., Huang, S., Mann, A. J., Waldmann, J., Weber, M., Klindworth, A., Otto, A., Lange, J., Bernhardt, J., Reinsch, C., Hecker, M., Peplies, J., Bockelmann, F. D., … Amann, R. (2012). Substrate-Controlled Succession of Marine Bacterioplankton Populations Induced by a Phytoplankton Bloom. Science, 336(6081), 608–611. 10.1126/science.1218344

Torruella, G., Grau-Bové, X., Moreira, D., Karpov, S. A., Burns, J. A., Sebé-Pedrós, A., Völcker, E., & López-García, P. (2018). Global transcriptome analysis of the aphelid Paraphelidium tribonemae supports the phagotrophic origin of fungi. Communications Biology, 1(1), 231. 10.1038/s42003-018-0235-z

Villaró, S., Sánchez-Zurano, A., Ciardi, M., Alarcón, F. J., Clagnan, E., Adani, F., Morillas-España, A., Álvarez, C., & Lafarga, T. (2022). Production of microalgae using pilot-scale thin-layer cascade photobioreactors: Effect of water type on biomass composition. Biomass and Bioenergy, 163, 106534. 10.1016/j.biombioe.2022.106534

Wang, X., & Hong, Y. (2022). Microalgae biofilm and bacteria symbiosis in nutrient removal and carbon fixation from wastewater: a review. Current Pollution Reports, 8(2), 128–146. 10.1007/s40726-022-00214-x

Waśkiewicz, A., & Irzykowska, L. (2014). Flavobacterium spp. – Characteristics, occurrence, and toxicity. In Encyclopedia of Food Microbiology (pp. 938–942). Elsevier. 10.1016/B978-0-12-384730-0.00126-9

Wei, Z., Wang, H., Li, X., Zhao, Q., Yin, Y., Xi, L., Ge, B., & Qin, S. (2020). Enhanced biomass and lipid production by co-cultivation of Chlorella vulgaris with Mesorhizobium sangaii under nitrogen limitation. Journal of Applied Phycology, 32(1), 233–242. 10.1007/s10811-019-01924-4

Wickham, H. (2020). Reshape v.2 1.4.4. 10.32614/CRAN.package.reshape2

Wickham, H. (2021). Tidyverse v.1.3.1. 10.32614/CRAN.package.tidyverse

Wickham, H., Chang, W., Henry, L., Pedersen, T. L., Takahashi, K., Wilke, C., Woo, K., Yutani, H., Dunnington, D., & Brand, T. van den. (2016). Ggplot2 v.3.5.1. 10.32614/CRAN.package.ggplot2

Wickham, H., François, R., Henry, L., Müller, K., Vaughan, D., & Software, P. (2023). Dplyr v.1.14. 10.32614/CRAN.package.dplyr

Wijffels, R. H., & Barbosa, M. J. (2010). An outlook onmMicroalgal biofuels. Science, 329(5993), 796–799. 10.1126/science.1189003

Zeileis, A., Grothendieck, G., Ryan, J. A., Ulrich, J. M., & Andrews, F. (2023). Zoo v.1.8–12. 10.32614/CRAN.package.zoo

Zhou, Q., Li, K., Jun, X., & Bo, L. (2009). Role and functions of beneficial microorganisms in sustainable aquaculture. Bioresource Technology, 100(16), 3780–3786. 10.1016/j.biortech.2008.12.037

