## Supplementary material for "The complex interplay between microalgae and the microbiome in production raceways": Method S1.docx

The PCR was performed in 25 μl volume with 0.08 μM primer concentration and NEBNext Q5 Hot Start HiFi PCR Master Mix (ref. M0543L, New England Biolabs). Cycling conditions were initial denaturation of 30 s at 98 °C followed by 5 cycles of 98 °C for 10 s, 48°C for 5 min, and 65 °C for 45 s. PCRs were treated with Exonuclease I (New England Biolabs) for 5’ at 37ºC and the enzyme was inactivated at 80ºC for 1’. After the first PCR, a second PCR was performed in a total volume of 50 μl. The reactions comprised NEBNext Q5 Hot Start HiFi PCR Master Mix and Nextera XT v2 adaptor primers. PCR was carried out to add full-length Nextera adapters: initial denaturation of 30 s at 98 °C followed by 17 cycles of 98 °C for 10 s, 48 °C for 30 s, and 65 °C for 45 s, ending with a final elongation step of 5 min at 65 °C.

Libraries were purified using AgenCourt AMPure XP beads (ref. A63882, Beckman Coulter) with a 0.9X ratio according to manufacturer’s instructions and were analyzed using Fragment Analyzer (ref. DNF-915, Agilent Biosystems) to estimate the quantity and check size distribution. Normalized libraries were prepared for subsequent sequencing.
