## Supplementary material for "The complex interplay between microalgae and the microbiome in production raceways": Method S2.docx

Categorization of the samples

- Samples with more than 0.7 relative abundance of *Desmodemus* *armatus* are categorize as healthy group.
- Samples between 0.1 and 0.2 relative abundance of *Desmodemus* *armatus* are categorize as unhealthy group.

Samples selected

| **TS** | **Reactor** | **nº healthy samples** | **nº unhealthy samples** |
| --- | --- | --- | --- |
| TS1 | WW | 16 | 14 |
| TS1 | FER | 17 | 22 |
| TS2 | WW | 13 | 17 |
| TS2 | FER | 22 | 14 |
| **ts1.ww.healthy.samples**: "PR173" "PR181" "PR189" "PR201" "PR205" "PR217" "PR221" "PR225" "PR233" "PR253" "PR254" "PR261" "PR265" "PR287" "PR531" "PR539" | | | |
| **ts1.ww.unhealthy.samples**:"PR84" "PR134" "PR154" "PR218" "PR222" "PR298" "PR384" "PR412" "PR424" "PR432" "PR498" "PR506" "PR520" "PR532 | | | |
| **ts1.fer.healthy.samples**:"PR421" "PR429" "PR435" "PR441" "PR449" "PR455" "PR463" "PR469" "PR475" "PR483" "PR489" "PR495" "PR503" "PR509" "PR523" "PR529" "PR537" | | | |
| **ts1.fer.unhealthy.samples**:"PR85" "PR93" "PR110" "PR113" "PR126" "PR136" "PR148" "PR172" "PR179" "PR184" "PR204" "PR212" "PR275" "PR285" "PR295" "PR303" "PR381" "PR430" "PR436" "PR510" "PR530" "PR538" | | | |
| **ts2.ww.healthy**.samples:"PR639" "PR647" "PR653" "PR659" "PR667" "PR673" "PR679" "PR687" "PR707" "PR713" "PR721" "PR727" "PR733" | | | |
| **ts2.ww.unhealthy**.samples:"PR648" "PR660" "PR748" "PR824" "PR862" "PR876" "PR910" "PR970" "PR1186" "PR1194" "PR1200" "PR1206" "PR1214" "PR1218" "PR1224" "PR1232" "PR1236" | | | |
| **ts2.fer.healthy samples**: "PR625" "PR631" "PR637" "PR645" "PR651" "PR657" "PR665" "PR671" "PR677" "PR960" "PR968" "PR984" "PR988" "PR1082" "PR1160" "PR1164" "PR1172" "PR1178" "PR1184" "PR1192" "PR1198" "PR1216" | | | |
| **ts2.fer.unhealthy.samples**:"PR652" "PR779" "PR840" "PR848" "PR854" "PR860" "PR894" "PR900" "PR908" "PR950" "PR1050" "PR1056" "PR1069" "PR1223" | | | |
