## Supplementary figures and images for "The complex interplay between microalgae and the microbiome in production raceways"

### Figure S1.pdf

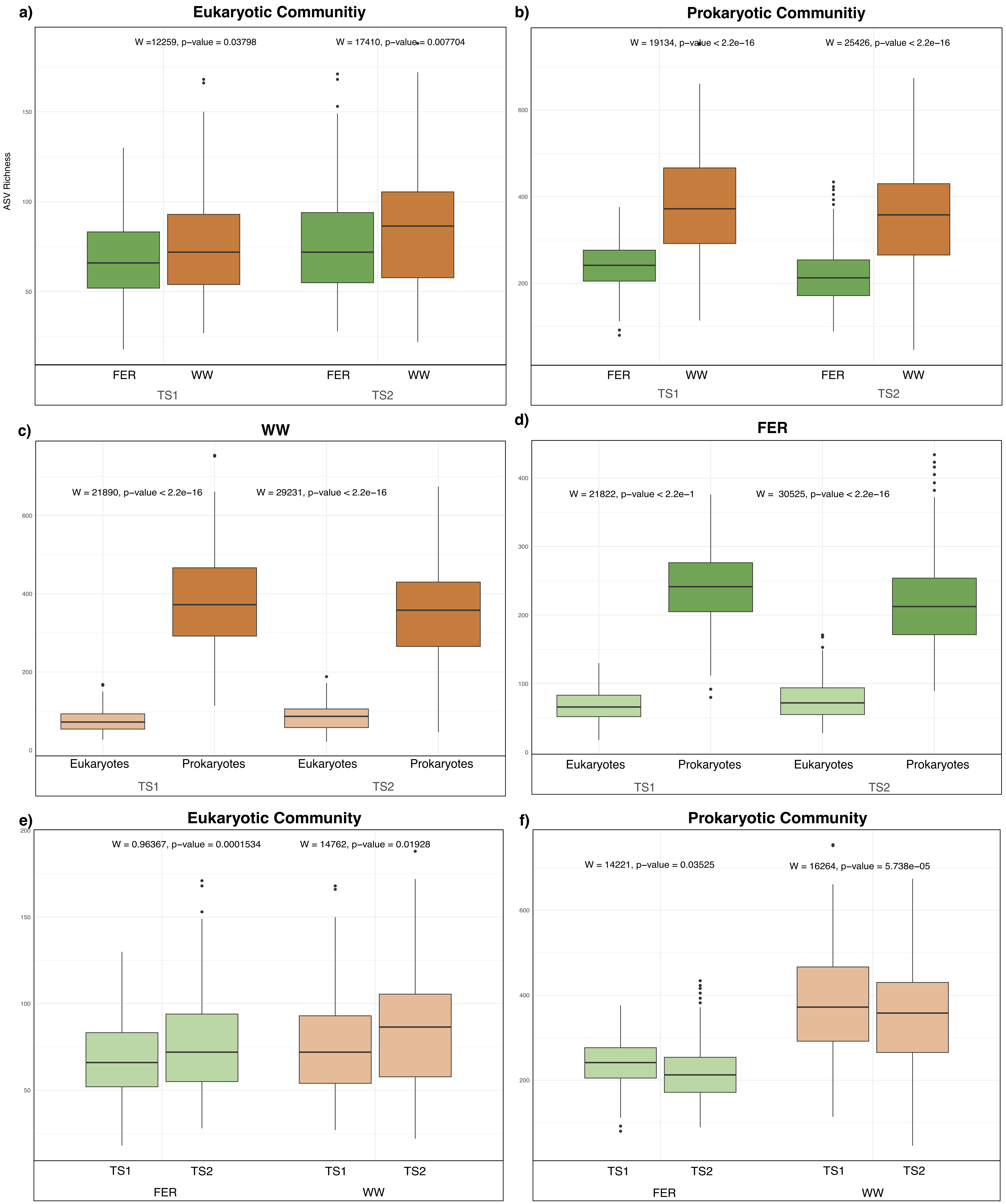

### Figure S2.pdf

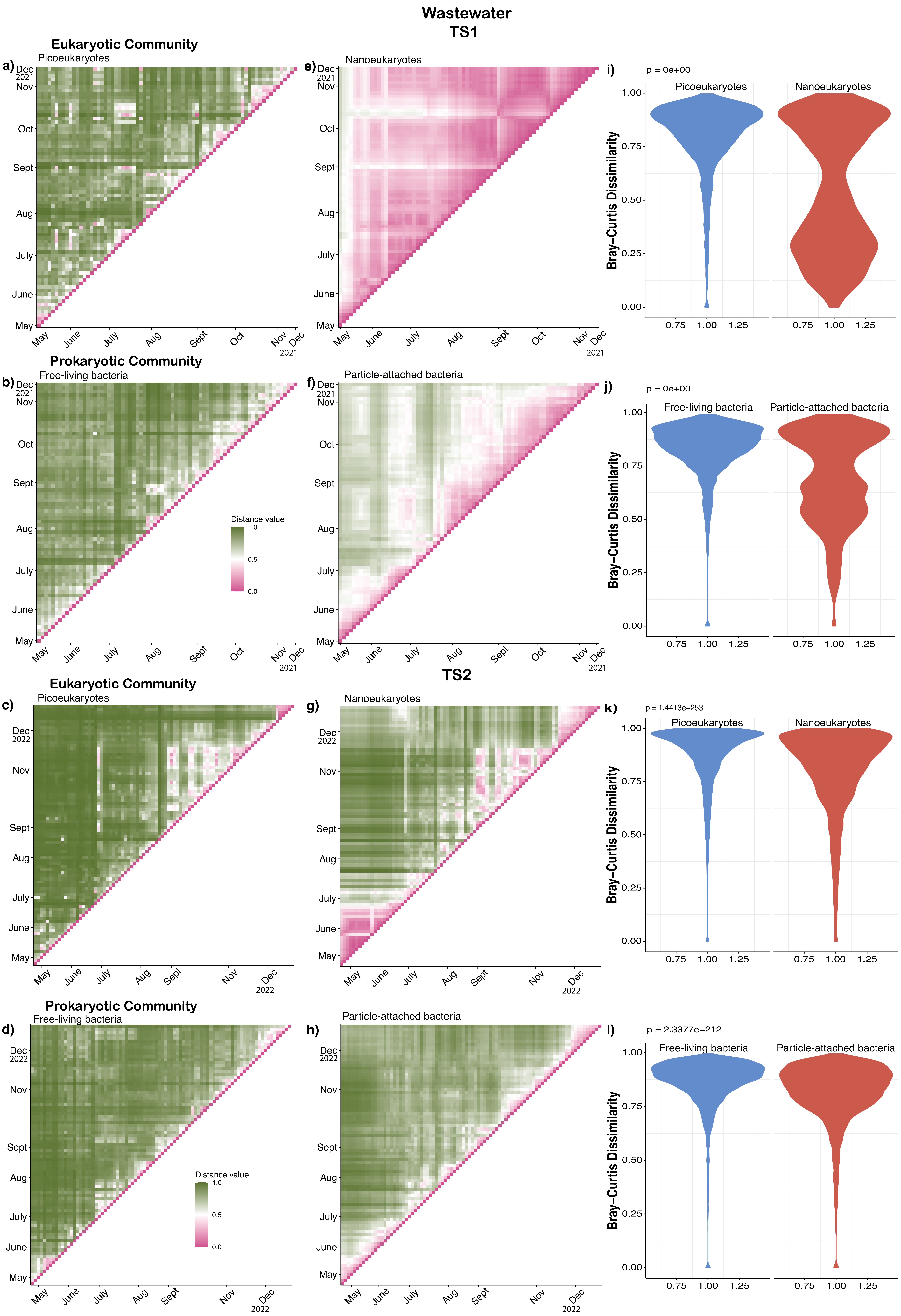

### Figure S3.pdf

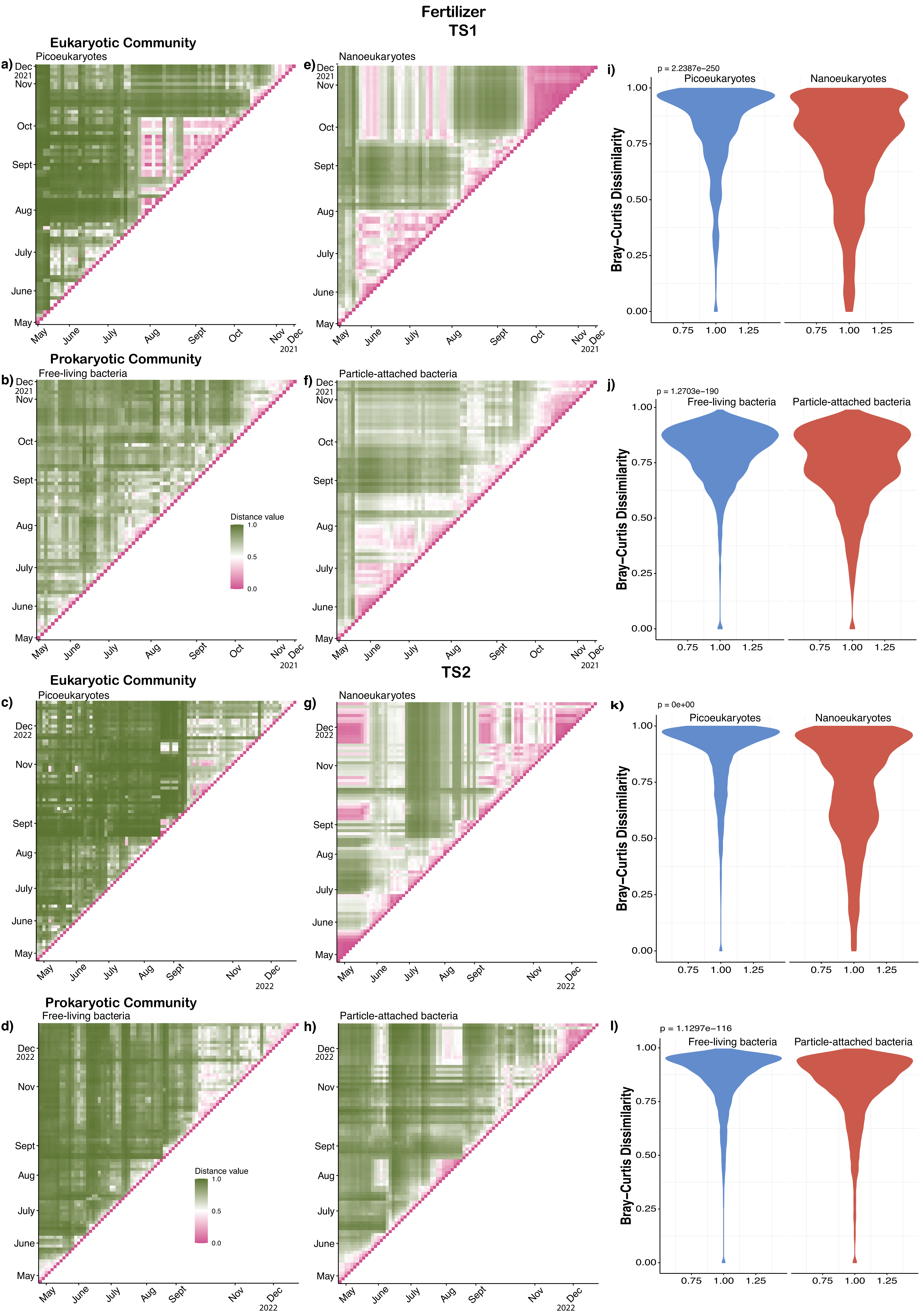
